# Stitching genomics data to protein structures: Virulence factors in non-O157 Shiga toxin-producing *Escherichia coli*

**DOI:** 10.64898/2026.01.09.698596

**Authors:** Sony Malhotra, Ashley Ward, Lauren Giles, Tom Gerrard, Martyn Winn, Nicola Holden

## Abstract

The genetic diversity of the Shiga toxin-producing *Escherichia coli* (STEC) sub-species group poses major challenges for assigning pathogenic potential, and for accurate diagnostics. Although existing risk-based frameworks work for the predominant serotypes e.g. STEC O157 or O26, they are less useful for more diverse genotypes, with variation in the genetic complement of virulence factors that is responsible for disease outcome. Moreover, genetic diversity occurs at the virulence factor allele level. We investigated the functional potential of genetic variation at a whole genome level and at an allele variant level, based on the premise that genomic variability underlies phenotypic traits including disease outcomes. We analysed 286 non-O157 STEC genomes for their virulence factor complement and determined co-occurrence patterns. Genetic variation in the well-characterized virulence factor intimin (*eae*) and its receptor (*tir*) was analyzed at a sequence level, and by modelling the three-dimensional protein–protein complex using AlphaFold3 and homology modelling. Different subtypes of Eae were shown to preserve their interaction with the receptor Tir by retaining the interactions at the protein-protein interface. The study shows that virulence factor characterisation in association with 3-D modelling of protein complexes can aid genomic analysis for risk assessment, and inform on functional mechanisms for pathogenicity.

## 1. Introduction

Shiga toxin-producing *Escherichia coli* (STEC) are a group of foodborne pathogenic bacteria that are predominantly transmitted to humans through contaminated food or from direct contact, particularly from farmed ruminant animals. Diseases caused by STEC range from diarrhoea (D), blood diarrhoea (BD) to more severe haemolytic uremic syndrome (HUS). Animals harbouring STEC do not typically demonstrate signs of infection, creating barriers to risk identification. Transmission of the pathogens through environmental routes, like water, creates a risk of infection from ready-to-eat produce. Treatment of infected patients is limited as antibiotic usage is (for the most part) contra-indicated due to increased risk of release of the Shiga toxin. As such, risk management with surveillance of multiple critical control points is the main means to prevent STEC transmission into the food chain.

Detection of the STEC group of pathogens is challenging due to their serological and genetic diversity. Serotypes other than O157 are increasingly detected and have become more prevalent^1,2^, with >338 detected in clinical infections^3^. In many European nations, serogroup O157 continues to be associated with the highest number of HUS cases, hospitalisation and bloody diarrhoea, followed by O26^4^. Identification of pathogenic non-O157 STEC is complicated as not all have the ability to cause disease^5^. As such, there is a pressing need for methods that can first identify them easily, and second, accurately indicate the likely level of risk.

Risk classification of STEC is currently based on the carriage of a particular set of virulence genes, as defined by a Joint FAO / WHO Expert Meeting on Microbiological Risk Assessment (JEMRA) Group^6^. The set of criteria and resulting decision tree (known as the JEMRA STEC risk matrix) are universally used as a means to classify pathogenicity, based on the Shiga toxin (*stx*) variant and carriage of adherence factors including initimin (*eae*) and / or Aggregative Adherence Fimbriae (*aaf*) adhesin (*aggR*). Hazard classification took into account other putative adherence factors, different serotypes and hybrid pathotypes of *E. coli*^6^. The JEMRA risk classification matrix can be supplemented by ranking risks associated with outcomes of clinical disease severity^4^ and using them as predictors for disease^7^, also based on *stx* variant and carriage of *eae*. The presence of *stx2* is strongly associated with severe disease progression, with highly enhanced potency compared to *stx1*^8^. Subclasses of *stx* can bind different glycosphingolipid receptors, although Gb3 is the principal entry point^9^. The *eae* gene is located on the 35-kb LEE^10^ (locus of enterocyte effacement) pathogenicity island along with other effectors and genes for the type 3 secretion system (T3SS). The presence of *eae* together with *stx*2 is commonly used as a molecular signal to predict pathogenic outcome in clinical settings^11,12^. Intimin (Eae) binds to the translocated intimin receptor (Tir), with the corresponding *tir* gene also located on the LEE. Tir is injected by the T3SS into host cells^13^ and embeds itself into the host membrane so that it can bind Eae^14^. For enterohaemorrhagic strains (EHEC *i.e.* most STEC), this is a precursor to the formation of attaching and effacing (A/E) lesions and actin-filament rearrangements resulting in pedestal formation^15^. Similarly in enteropathogenic *E. coli* (EPEC, a different diarrheagenic *E. coli* that is also characterised by A/E lesions but does not encode the Shiga toxin gene), *eae-tir* also facilitates intimate host binding^16^.

It is notable that a relatively high proportion of hospitalisation and bloody diarrhoea disease occur in the absence of *eae* carriage^4^. As the minimum virulence gene repertoire is still unclear, computational methods based on genomic content are now being tested to help with this determination. Machine learning approaches have predicted that serious disease (BD and HUS) is associated with additional factors in the O157 serotype beyond established virulence factors^17^.

The numbers of STEC serotypes other than O157 is increasing year-on-year across the UK, and in some regions has become more dominant. In England, there were 2,762 total cases of non-O157 STEC reported in 2024, an increase of 22% from 2023 and 175% compared to 2019^1^. In contrast, there were 564 confirmed cases of STEC serotype O157 in England (2024^1^), a decrease of 26% compared to 2022^18^. The same trend is seen in Scotland with increasing non-O157 STEC, of 182 laboratory reports in 2023^2^ compared to 170 in 2022^18^. The level of STEC serotype O157 is proportionally lower in Scotland, with 133 laboratory reports in 2023, an decrease from a spike of 270 reported in 2022 (PHS GIZ 2020-2021 report^19^). In European member states there has been a continuous increase in serogroups O26, O146, O145, and O103, although O157 remains the most frequently reported (19%)^20^. The main non-O157 STEC serotypes that are associated with clinical disease include O26:H11 and O145:H28 (as listed in Enterobase^21^, 1952-2024, Table S1). Genomic diversity that occurs between serotypes encompasses surface factors affecting attachment phenotypes, virulence factors affecting pathogenicity and host colonisation, and horizontally acquired elements including prophage, insertion elements and plasmids^22^.

Genomic variation is a reflection of selection pressure on the gene products and corresponding protein function. This occurs where a genomic complement or gene allele type has been under positive selection in a given environment or host, and so represents the successful clone. In turn, it raises the possibility that such genomic variation can be used to indicate particular traits as a result of host or habitat niche adaptation, such as pathogenicity within a human host or colonisation of secondary hosts. This broad principle has been applied to another set of pathogens, *Salmonella* species, where machine learning has been applied to protein functional domains to assign biomarkers related to disease severity potential^23^.

To investigate the pathogenic potential of less well characterised non-O157 STEC, a set of clinical isolates (n=286) was collated and the key virulence factors identified through genome analysis. A reference panel of known virulence factors^24,25^ was used to map these isolates for the presence and distribution of the corresponding genes. Co-occurrence patterns of these genes were then assessed to explore possible functional relationships that may dictate their pathogenic potential. As expected, the *eae* and *tir* virulence genes, associated with A/E lesions, were consistently associated, indicating the presence of the LEE pathogenicity island in a subset of isolates^26^. Within this subset, we could identify multiple eae subtypes and further allelic variation within the subtypes.

We propose that the genomic variability underlying such traits can be used in diagnostics applications. Therefore, investigating allelic variations can help understand their impact on potential protein function and pathogenicity. Three-dimensional protein structure modelling (using AlphaFold3^27^) was used to model the Intimin-Tir complexes and map the allelic variations onto the structures for assessing the impacts on protein structure and function.

## 2. Materials and Methods

### 2.1 Data Collection

Human (clinical) isolates were selected from Enterobase^21^ (queried Dec 2024), using the following filters:

a. Geographical location: United Kingdom; UK; Great Britain; GB; England; Scotland; Wales; Northern Ireland /NI (but not ‘Ireland’)
b. Source type: Human
c. ‘*E. coli’* only and Species: exclude all isolates including “*Shigella*”
d. Serotype filter: anything with O and H type included, all other descriptors removed. Included ones with O type only, removed all blanks or listed as ‘Escherichia coli’. All the O157 were removed as the focus of this study is non-O157.
e. Path/Non path: Pathogenic & blanks included
f. Pathotypes related to UTI or ExPEC removed. Included EIEC.
g. Lab contact: Anything that contain ‘Public Health England’ included in email addresses
h. Collection year: from 2010 onwards

These entries in Enterobase do not have clinical outcome data because questionnaires are not routinely administered to non-O157 cases in the same way as they are STEC O157 cases (pers. communication Claire Jenkins).

Within the non-O157 STEC dataset, O26 STEC isolates were eliminated as it is the most prevalent non-O157 STEC serotype in the UK^1,2^ and could disproportionately skew the dataset and subsequent analyses. Furthermore, O26 is already one of the most extensively characterized non-O157 serotypes. The aim was to obtain a broader snapshot of recently emerged and less-characterized strains. After applying the filters described above and removing O26, to further capture H-antigen diversity and maximise the diversity of serotypes, an isolate per H-antigen type was selected for each non-O157 serotype.

Further, these isolates were filtered for the presence of *stx* gene to identify the (non-O157) STEC genomes. For this, the paired end reads from Sequence Read Archive^28^ (SRA) were downloaded using the fastq-dump utility, and filtered using the BBTools^29^. The following criteria were used for cleaning the downloaded reads:

i. discard reads with average quality below 20.
ii. Trim adapters based on pair overlap detection (does not require known adapter) using BBMerge
iii. Trim both reads to the same length
iv. Also used the generalized adapters and trimmed provided by BBTools package which includes Illumina Truseq and Nextera adapter sequences

The obtained cleaned reads were then mapped to the *E. coli* stx gene sequences downloaded from the VirulenceFinder 2.0 (as described in section 2.2), using bowtie2^30^ and samtools^31^. The downloaded set of stx sequences from VirulenceFinder 2.0 cover a variety of stx subtypes and include subtypes stx1a, stx1c, stx1d, stx1e, stx2a, stx2b, stx2c, stx2d, stx2e, stx2f, stx2g, stx2h, stx2i, stx2j, stx2k, stx2l, stx2m, stx2n and stx2o.

The resulting stx+ genomes were assembled using the Shovill pipeline (https://github.com/tseemann/shovill) and the Spades assembler^32^ which scans over multiple kmer lengths. Prokka^33^ was used to annotate the assembled genomes and identify genes.

### 2.2 Virulence Factors Mapping

The list of *E. coli* virulence factors (virulence_ecoli.fsa) was downloaded from the Virulence Finder 2.0^24,25^ (database version 2022, last updated in April 2024, https://bitbucket.org/genomicepidemiology/virulencefinder_db/src/master/). For some of the virulence factors such as intimin (*eae*), the downloaded annotations include the subtype mapping. The whole list comprises 4942 sequences of virulence factors including the subtypes and their multiple sequences, which maps to 680 sequences for different subtypes of virulence factors by removing alleles.

Virulence Finder provides a very broad list that includes any factor associated with pathogenicity, but many are simply colonisation factors and not true virulence genes. The sequence identity between related entries can also be high. Thus, *stx1* and *stx2* subtypes are just a few SNPs different and there can be multiple subtypes per genome which may cause the assembly to split the gene.

An in-house database was built using BLAST^34^ for the reference virulence factor sequences which was then used to search for virulence factors in the Prokka-annotated isolate genomes. tblastn^35^ was used to query the database with an evalue cut off 10^-4^. The hits were then filtered using the following criteria:

1. percent identity >=90%
2. query coverage >= 75%, where query coverage is the aligned query/length of query

Using these two filters if a sequence maps to more than one virulence factor gene, the hit with the greatest percent identity was selected. Further filtering was done using the subject (virulence factor gene) coverage cut off of >=75%, where subject coverage is the aligned subject/subject length to filter out the genome sequences which are just partial sequences.

For isolates carrying the intimin gene, the main genetic subtype (alpha, beta, gamma, etc) was identified based on the Virulence Finder annotation of the closest sequence match, noting that the database includes several entries for each subtype. Although Virulence Finder contains sequences for the intimin receptor, encoded by the *tir* gene, there is no equivalent subtype annotation for these entries.

The term ‘variant’ is used throughout the text and is defined in context, with respect to STEC serotype, gene presence/absence at the genome level, specific gene subtypes, or within gene point mutations.

### 2.3 Co-occurrence of virulence factors

Entries in a co-occurrence matrix were defined as the number of isolates in which a pair of virulence factors occur together, normalised by the total occurrence of the virulence factor on the row, so that the matrix is asymmetric with respect to the two virulence factors. An enrichment factor was also defined of a virulence factor *A* with respect to a virulence factor *B* as the ratio of the occurrence of *A* in *B*+ isolates to its occurrence in *B*- isolates, adjusted for the relative number of each group of isolates:

Enrichment of virA in presence of virB: E_A_ ^(B)^ = (virAB/virB) / ((virA – virAB) / (286 – virB))

A value greater than one implies that a virulence factor *A* is more likely to be found in an isolate containing *B* than one where *B* is absent. The Jaccard similarity was also considered, J_AB_, defined as the number of isolates containing both virulence factors divided by the number of isolates that contain at least one.

### 2.4 Sequence alignment and phylogenetic trees

Sequence alignments were performed using MUSCLE^36^. RaxML^37^ was used to generate phylogenetic trees for different sets of sequences. For extracting the LEE island, the LEE sequence from the *E. coli* O157:H7 strain Sakai (GenBank accession no. BA000007) was used. BLAST^34^ against the Sakai sequence was performed using all the contigs from the *eae*+ genomes. A contig was considered to contain a LEE island if it shared at least 80% sequence identity with the Sakai LEE sequence and the alignment length was at least 20 kb. The LEE sequences were extracted from the *eae*+ isolates (n=85) and the pangenome was built using Panaroo^38^.

The pangenome for the *stx+* isolates (n=286) was built using Panaroo^38^ and the annotation previously output from Prokka. RaxML (v8.2.12)^37^ was used to build an alignment of the core genome. Visualisation of the annotated trees was done in Iroki^39^.

Sequence alignments were used as an input to SplitsTree^40^ to compute a split decomposition network (selecting network -> split decomposition).

### 2.5 Three dimensional structural modelling of virulence factors

AlphaFold3^27^ was used to model the structures of protein complexes. The protein sequences of Eae (D1, D2 and D3 domains) and Tir (Tir intimin binding domain) in Fasta format were used as input to the AlphaFold3 server (https://alphafoldserver.com/). In total, ten subtypes of Eae were modelled using sequences taken from isolates for which the subtype had been identified, with the cognate Tir sequence taken from the same isolate. The following accession IDs were used for structural modelling of Eae subtypes: alpha:SRR3581427, beta:SRR4180889, gamma:SRR3581353, rho:SRR3241980, theta:SRR4179996, epsilon:SRR3574270, xi:SRR16203499, delta:SRR29176816, iota:SRR4195732 and zeta:SRR4184837. For each subtype, a separate job was run, using the Eae protein sequence as entity 1 (copy = 1) and the Tir protein sequence from the same genome as entity 2 (copy = 1). The server returns top five scoring complexes and the one with highest ranking_score was selected. Ranking_score combines pTM (predicted TM-score for full structure), ipTM (predicted interface TM-score), penalises clashes and encourages the absence of spurious helices in disordered regions^27^.

Additional homology based structural models were built for some of the subtypes (rho and theta) using Modeller (v9.10)^41^. The template used for the homology modelling is PDB ID: 1F02 and the model with lowest DOPE (Discrete Optimised Potential Energy) score was selected as the best model.

## 3. Results

### 3.1 Collation of STEC isolate genomes

#### 3.1.1 Initial download

A retrieval of all E. coli isolates from EnteroBase yielded 86,609 accessions. The filtering criteria described in Section 2.1 (Methods) were applied to retain clinical, non-O157 isolates, resulting in 7,855 isolates. The most common serotypes (Figure 1A) in this subset were O26 (n = 1,027), O145 (n = 554), and O146 (n = 515). The O26 serotype was subsequently excluded because it is already well characterised and overrepresented in public databases. To ensure a broader snapshot of recently emerged and less-characterized strains, one isolate per H-antigen was retained for each remaining non-O157 serotype. This resulted in a filtered subset of 1177 isolates.

**Figure 1:**
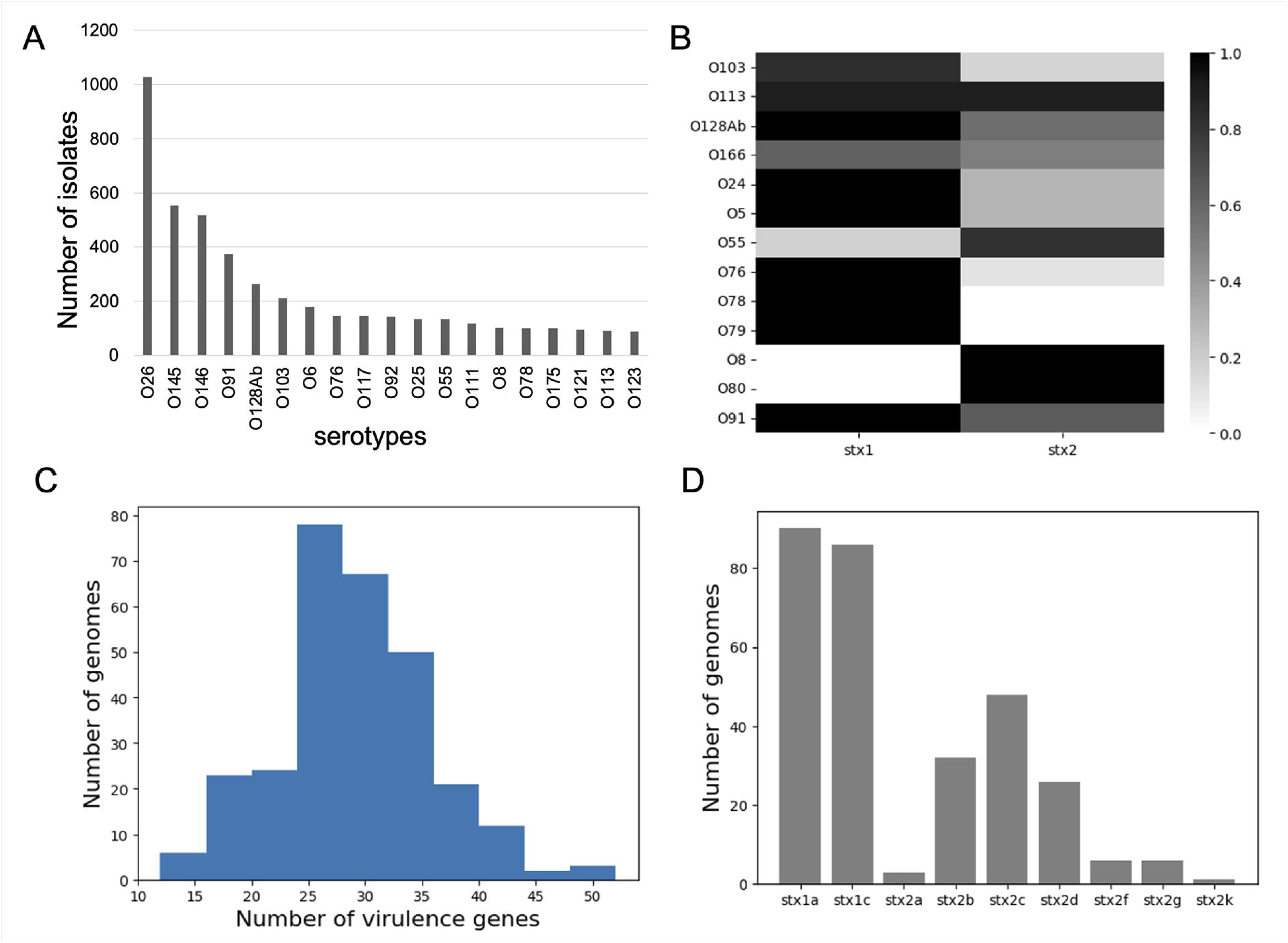
Collection of non-O157 STEC isolates (n=286) and virulence factor mapping. **A**. Distribution of different serotypes in the collection of non-O157 STEC genomes from the clinical isolates obtained from the Enterobase. **B**. Presence of different types of *stx* in these serotypes. **C**. Histogram of number of virulence factors in non-O157 STEC genomes. **D**. Subtypes of *stx* toxin genes in non-O157 STEC genomes.

Entries in Enterobase do not have clear metadata on pathogenicity or clinical outcome. We make the assumption that all of the human isolate sequences selected from Enterobase using this protocol were associated with clinical disease symptoms, since the isolates in this collection were all deposited by reference or diagnostic laboratories. They are hereafter referred to as ‘clinical isolates’.

#### 3.1.2 Presence of Shiga toxin gene

Isolate genomes could not be filtered in Enterobase by the presence of *stx* gene(s) because of incomplete metadata, nor were they generally designated explicitly as STEC. Therefore, the genome sequence reads were downloaded and mapped to the *E. coli stx* gene alleles from the Virulence Finder v2.0 database^24,25^. Out of the 1177 genomes in the original Enterobase set, 326 genomes had at least one read mapped to one of the *stx* gene subtypes. This does not necessarily imply that all these 326 genomes are true STEC due to the low threshold of number of reads mapped to the *stx* genes. Therefore, confirmation of *stx* presence was carried out at the annotation level following assembly of the 326 genomes.

#### 3.1.3 Filtering for STEC genomes

The 326 putative STEC genomes were assembled and annotated, as described in Section 2.1. A more stringent check for the presence of *stx* genes was based on *tblastn* searches, using the protein sequences obtained after the annotation step (section 2.2). As the reference *stx* genes (from Virulence Finder v2.0) includes both subunit A and subunit B genes, the BLAST hits were further validated, and only genomes that included both *stx* toxin A and B subunits were selected. This resulted in 286 *stx+* isolates, henceforth referred to as non-O157 STEC isolates. The Enterobase accession numbers for these 286 isolates are provided in Table S2. The distribution of *stx* types among serotypes present in more than five isolates is shown in Figure 1B. Sole carriage of *stx2* only occurred in serotype O8 and O80, while conversely sole carriage of *stx1* occurred in O78 and O79 serotypes. All other serotypes (9) encoded both *stx1* and *stx2* subtype reads.

### 3.2 Virulence factor genes in STEC genomes

#### 3.2.1 Virulence factors

The complete set of virulence factor alleles was mapped onto non-O157 STEC genomes (n=286) using tblastn^35^ following the annotation of the assembled genomes (see section 2.2). For ease of presentation, virulence factors are referred to by their 3-letter gene name, of which there were 85 present in the set of non-O157 STEC isolates (Table S3) . Where appropriate, individual genes were identified, of which there were 130 in the genome set when allelic variants were merged (Table S4). Isolate genomes varied in the number of virulence factors identified, and Figure 1C shows the distribution of virulence factor load. The genome with maximum number of virulence factors is accession SRR3579389 with 52 factors mapped to it, in contrast to minimum number of virulence factors (n=12) in the genome of accession SRR29241024.

For the two main *stx* groups, 128 genomes encoded only the *stx1* subtype, 110 had only *stx2* subtype, and the remaining 48 genomes contained representatives of both *stx1* and *stx2*. Further division into *stx* subtypes was carried out using the reference sequences in Virulence Finder v2.0 (Figure 1D). We observed two subtypes of *stx1* (1a and 1c, roughly in equal numbers) and seven subtypes of *stx2* (2a,2b,2c,2d,2f,2g,2k) across the 286 isolates (Figure 1D). Of these seven stx2 subtypes, the most common were *stx2b* and *stx2c*, with low numbers of five other subtypes.

The most frequent virulence factors were assessed, classed as present in at least 50% of the genomes (Figure 2A). *nlpI* (lipoprotein NlpI precursor) and *terC* (tellurium ion resistance protein) are present in all 286 genomes, followed by *yeh* (4 genes, YHD fimbrial cluster) and *csgA* (curli major subunit) which are present in 283 and 281 isolates respectively. Nine other virulence factors are widespread in our set of isolates, though still absent from a significant fraction.

**Figure 2:**
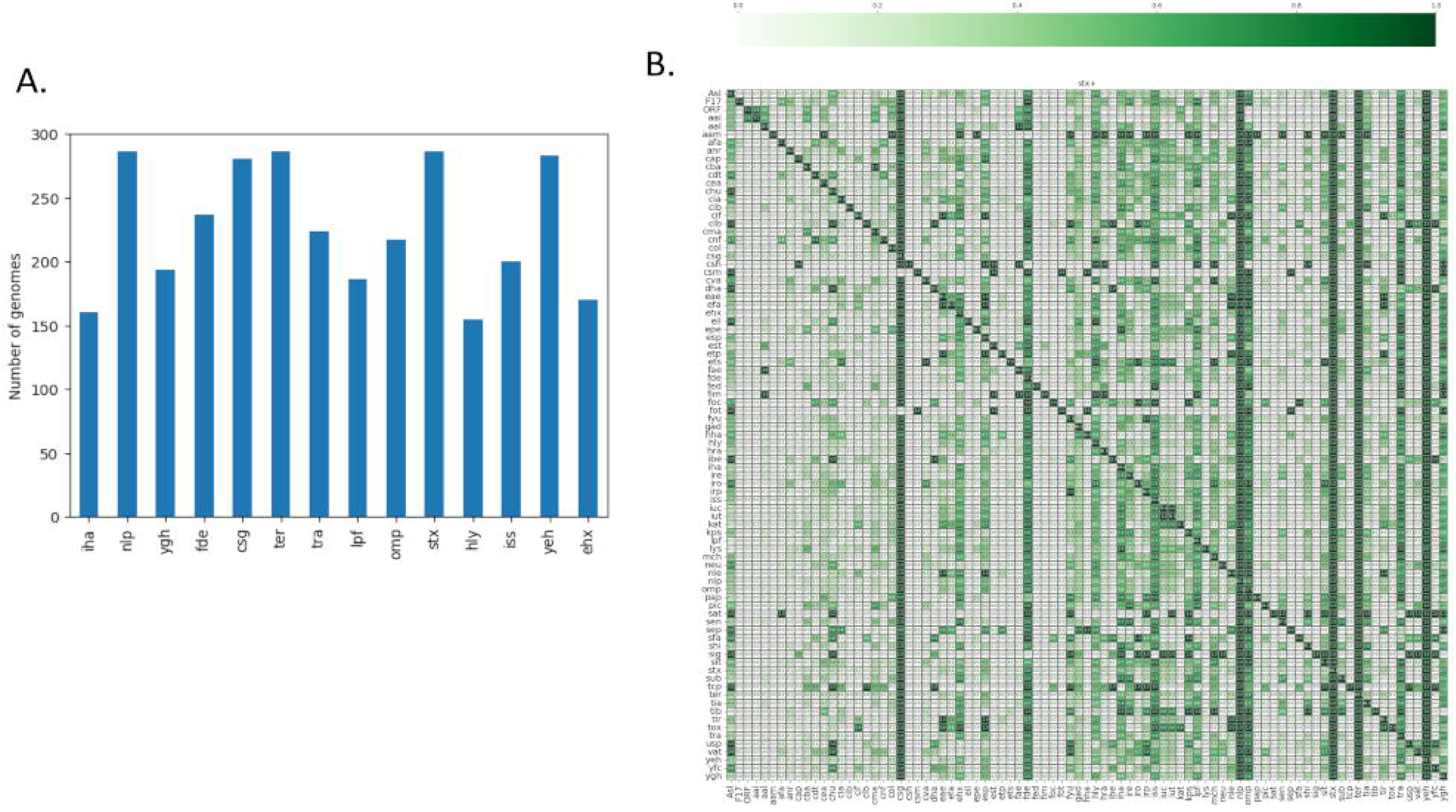
Virulence factors presence and co-occurence in the set of isolates. **A.** Most frequent virulence factors (present in at least 50% of the genomes). **B.** Co-occurrence of virulence factors in our set of 281 isolates. The diagonal gives the absolute number of isolates that a virulence factor occurs in. The off-diagonal entries give the number of isolates containing both virulence factors (corresponding to the row and the column), given as a fraction of the total number of isolates that the virulence factor on the row occurs in. Note that this definition means that the matrix is asymmetric.

Co-occurrence of virulence factors was determined for the set of 286 non-O157 STEC genomes (Figure 2B). The co-occurrence values were normalised for a pair of virulence factors by the total occurrence of the virulence factor on the row, so that the matrix is asymmetric with respect to the two virulence factors.

First, co-occurrence of virulence factors with intimin was a point of focus, as it is one of the key virulence factors for the STEC group^41^ in terms of incidence and disease outcome (see row ‘eae’ in Figure 2B) ^42^. To help identify significant correlations with intimin, an enrichment factor E_A_ ^(eae)^ was defined as the ratio of the occurrence of a virulence factor in *eae*+ isolates to its occurrence in *eae*- isolates, adjusted for the relative number of each group of isolates. The Jaccard similarity, J_A,eae_, between the set of isolates containing a specified virulence factor and the set carrying *eae* was also considered. Values for all virulence factors are listed in Tables S3 and S4. By these definitions (see Methods), *tir* has E_tir_^(eae)^ of infinity as it never occurs without its binding partner, and J_tir,eae_ = 1 since it has an identical distribution across the isolates. On the other hand, *etp* has E_etp_^(eae)^ of infinity as it only occurs in *eae*+ isolates, but J_etp,eae_ = 0.27 since it only occurs in 23 of the *eae+* isolates and thus has a different distribution.

The gene group *nleA, nleB, nleC* comprises non-LEE effectors which are functionally dependent on the LEE-encoded T3SS. At least one *nle* gene is found in 83 out of 85 *eae*+ isolates, while all three are found in a single *eae-* isolate. The presence of *eae* and *nle* genes is reported to be a signature for highly virulent STEC strains and are reported to co-occur in non-O157 serotypes^43^. *nle* is reported to correlate with outbreak and HUS potential^44^.

For the *cif* gene, 37 out of 38 occurrences are in *eae*+ isolates, while for both *espJ* and *efa1* it is 44 out of 45. Several of these genes have been observed on a lambdoid prophage ΦE22^45^ and so may be carried together. A single isolate (AIPCOOIH) carries the only occurrences of *nleA, nleB, nleC, cif, espJ* and *efa1* that are not associated with intimin.

Other genes enriched in *eae*+ isolates include *espP*, *tox*, *katP* and the bacteriocin *cia*. Table 1 lists other virulence factors that were observed to co-occur with eae, though some are also widely seen in *eae*- isolates. Some of these are toxins (such as hemolysin, enterohemolysin), adherence and colonisation factors (*efa, fdeC*), or help in evading the bacterial complement system (*iss*).

**Table 1:** Virulence factor genes that show significant co-occurrence with *eae* in non-O157 STEC isolates. The co-occurrence is defined as the number of isolates carrying both the virulence factor of interest and *eae*, as a fraction of the total number of *eae*+ isolates, corresponding to the values on the *eae* row of Figure 2B. The following column gives the absolute number of isolates carrying the virulence factor, divided between *eae*+ and *eae*-isolates.

| Gene co-occurring with *eae* | Co-occurrence | Number in *eae*<sup>+</sup> / *eae*<sup>-</sup> isolates | Short description |
| --- | --- | --- | --- |
| *cif* | 0.44 | 37 / 1 | Type III secreted effector |
| *efa1* | 0.52 | 44 / 1 | EC factor for adherence; lymphostatin *Efa1-LifA* |
| *ehxA* | 0.76 | 65 / 105 | Enterohaemolysin:known to be associated with HUS and diarrheal disease |
| *espA*, *espB*, *espC*, *espF*, *espI*, *espJ*, *espP*, *espY* | 1.00 | 85 / 48 | E.coli secreted protein; *espA*, *espB*, *espF* carried on LEE island; *espP* serine protease |
| *fdeC* | 0.73 | 62 / 175 | intimin-like adhesin *fdeC*: biofilm formation; structural similarity to intimin |
| *hlyA*, *hlyE*, *hlyF* | 0.67 | 57 / 98 | Hemolysin; known to be associated with HUS and diarrheal disease |
| *iha* | 0.46 | 39 / 121 | *irgA* homologue adhesin (*Iha*) |
| *iss* | 0.69 | 59 / 141 | Increased serum survival; complement resistance |
| *lpfA* | 0.45 | 38 / 148 | Long polar fimbriae: adhesive factor;also known to be expressed in LEE-negative clinical isolates |
| *nleA*, *nleB*, *nleC* | 0.98 | 83 / 1 | Non-LEE effectors |
| *ompT* | 0.94 | 80 / 137 | Outer membrane protease (protein protease 7): |
| *traJ*, *traT* | 0.68 | 58 / 166 | *traJ* (Positive regulator of conjugal transfer operon)) and *traT* (Outer membrane protein complement resistance:); |
| *yghJ* | 0.73 | 62 / 132 | Secreted and surface associated lipoprotein; aids in colonisation |

In contrast, many pairs of virulence factors are mutually exclusive, meaning that there are no isolates that contain both. This may simply be because of low overall occurrence in the dataset, or may reflect some aspect of genomic structure *e.g.* competition for the same phage integration site. In particular, the genes *subA*, *kpsM*, *pic* and *senB* occur 79, 92, 24 and 67 times respectively, but never in *eae+* isolates. The *subA* gene encodes a component of the SubAB toxin, which has been observed in LEE-negative STEC strains including clinical human isolates^46^. The gene *pic* encodes for a member of the serine protease auto-transporter (SPATE) family and has been shown to have mucolytic activity^47^.

The *ireA* gene occurs in 99 eae- isolates and a single *eae*+ isolate (SRR3579389). Similarly, *aal* genes occur in 28 *eae-* isolates and *fae* genes occur in 29 *eae-* isolates, while a single *eae+* isolate (SRR3573434) contains a copy of *aalF, aalH, faeE, faeF, faeC, faeI, faeD*.

Next, co-occurrence of groups of genes independent of intimin was determined. A co-occurrence matrix for a set of 32 genes was generated (Figure S1). Genes that occur either too rarely or which are very common were excluded, as were those that showed no significant co-occurrence. Several sets of genes were identified that showed a degree of co-occurrence between each other (Figure S1). The largest group consisted of *’ireA’, ‘kpsM’, ‘mchC’, ‘mchF’, ‘senB’, ‘shiA’, ‘sitA’, ‘subA’, ‘tia’*. These are all enriched in *eae-* isolates, and include three of the genes (*subA*, *kpsM* and *senB*) that are never found in *eae+* isolates. Similarly, adhesion-associated locus genes (*aal*) and fimbrial genes (*fae*) are found to co-occur, and are found more frequently in *eae-* isolates.

Another significant group consists of genes *cif*, *efa1*, *iucC*, *iutA*, *katP*. Three of these were observed above to co-occur with intimin, and therefore will co-occur with each other. Finally, there are pairs of genes which occur together, e.g. the Colicins B and M (*cba*, *cma*), as has been seen previously^47^, which are almost always found adjacently on the same large conjugative plasmid. The genes *cba* and *cma* have intimin enrichment factors of 0.26 and 0.43 suggesting a slight enrichment in *eae-* isolates.

#### 3.2.2 Intimin (*eae*) subtype mapping

Within the set of 85 (30%) isolates that are both *eae* and *stx* positive, several *eae* genetic subtypes can be distinguished (based on their best match upon BLAST searches against the reference *eae* sequences, section 2.2). The *E. coli* virulence factors in the virulence finder database (section 2.2) include 20 subtypes of *eae*: alpha, beta, delta, epsilon, eta, gamma, iota, kappa, lambda, mu, nu, omicron, pi, rho, sigma, tau, theta, upsilon, xi and zeta. Of these, 10 *eae* subtypes were identified in the set of non-O157 *stx*+ isolates; namely alpha, beta, delta, epsilon, gamma, iota, rho, theta, xi and zeta.

The *eae+* genomes were mapped to the *stx* gene subtypes in order to investigate if any *eae* subtype occurrence correlates with *stx* gene type (Figure 3A). 6/85 eae+ isolates possess both *stx1* and *stx2*, while 44 and 35 possess only *stx2* or *stx1*, respectively. Four *stx* gene subtypes, namely *stx1a, stx2c, stx2d* and *stx2f*, co-exist with the *eae* gene. *stx2* has been shown to be associated with severe disease phenotype^49,50^ and it is interesting to note that in the non-O157 STEC dataset, *eae*-gamma and *eae*-iota co-occur exclusively with the *stx2c* subtype. stx2c is associated with six further *eae* subtypes, although in those cases not exclusively. *stx2d* which is reported to be very virulent in mice and associated with HUS^51^ was seen co-occurring mainly with subtype *eae*-xi, but also with *eae*-epsilon. The less virulent form *stx1a* was present with 6 *eae* subtypes.

**Figure 3:**
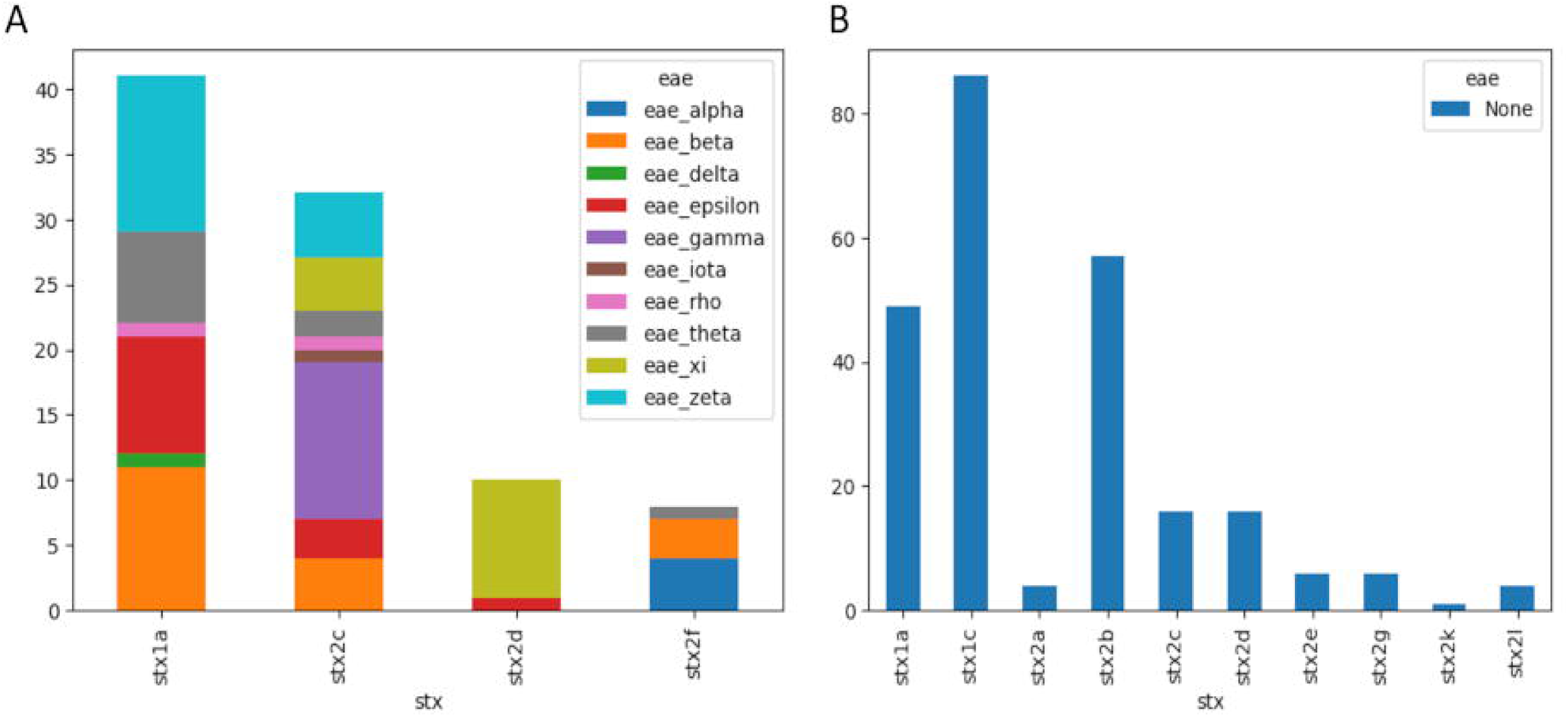
Co-occurrence of *stx* and *eae* genes. **A**. Distribution of intimin subtypes in *stx* subtypes. **B**. Distribution of *stx* subtypes in intimin negatives isolates.

The *stx* subtype mapping for *eae-* isolates (n=201) shows that 21% (n=42) of the isolates possess both *stx1* and *stx2*, while 33% (n=93) and 46% (n=66) of the isolates possess only *stx1* or *stx2* respectively. Further, mapping these to *stx1* and *stx2* subtypes, it is interesting to note that some subtypes (*stx1c, stx2a, stx2b, stx2e, stx2g, stx2k and stx2l*) were exclusively seen in the *eae-* set (Figure 3B).

#### 3.2.3 Evolution of the *eae* (intimin) gene

Across the set of 85 *stx*+/*eae*+ isolates, there are 31 unique translated intimin sequences, categorised into 10 subtypes. The diversity across the observed intimin sequences is displayed as a heatmap (Figure S2) generated using percent identity values between the protein sequences. The minimum percent identity is between the single iota sequence and two zeta subtype sequences (79.1%) whereas epsilon and xi sequences are most similar with ∼97% identity (Figure S2). To understand the evolution of the *eae* gene and the variation within the different subtypes, the intimin subtypes were mapped onto phylogenetic trees. As intimin is encoded by a gene on the LEE island, originally acquired through horizontal gene transfer, we sought to separate different evolutionary drivers by constructing phylogenies at the level of the core genome of *stx*+ isolates (n=286), the whole LEE island of *stx*+/*eae*+ isolates (n=85) and the individual translated intimin sequences (n=31).

The selected set of non-O157 STEC isolates (n=286) were observed to have a core genome (99% <= strains <= 100%) of only 3119 genes out of the total ∼36k genes, with most genes in the shell and cloud regions. This agrees with the known observation of pathogenic *E. coli* genomes to be larger and more diverse than their commensal counterparts^52^. Virulence factors such as *eae*/*tir* are assigned to the cloud region, in accordance with the low level of occurrence (highlighted above). A phylogeny was obtained using the core genome of the isolates, and annotated with the *stx* and *eae* subtypes (Figure 4A). The distribution of *eae* across different clades is consistent with multiple acquisitions of the LEE island via horizontal gene transfer. The phylogeny shows reasonable clustering of the *eae* subtypes, although several subtypes are split across different branches which may be due to inaccuracies in the phylogeny or from recombination events.

**Figure 4:**
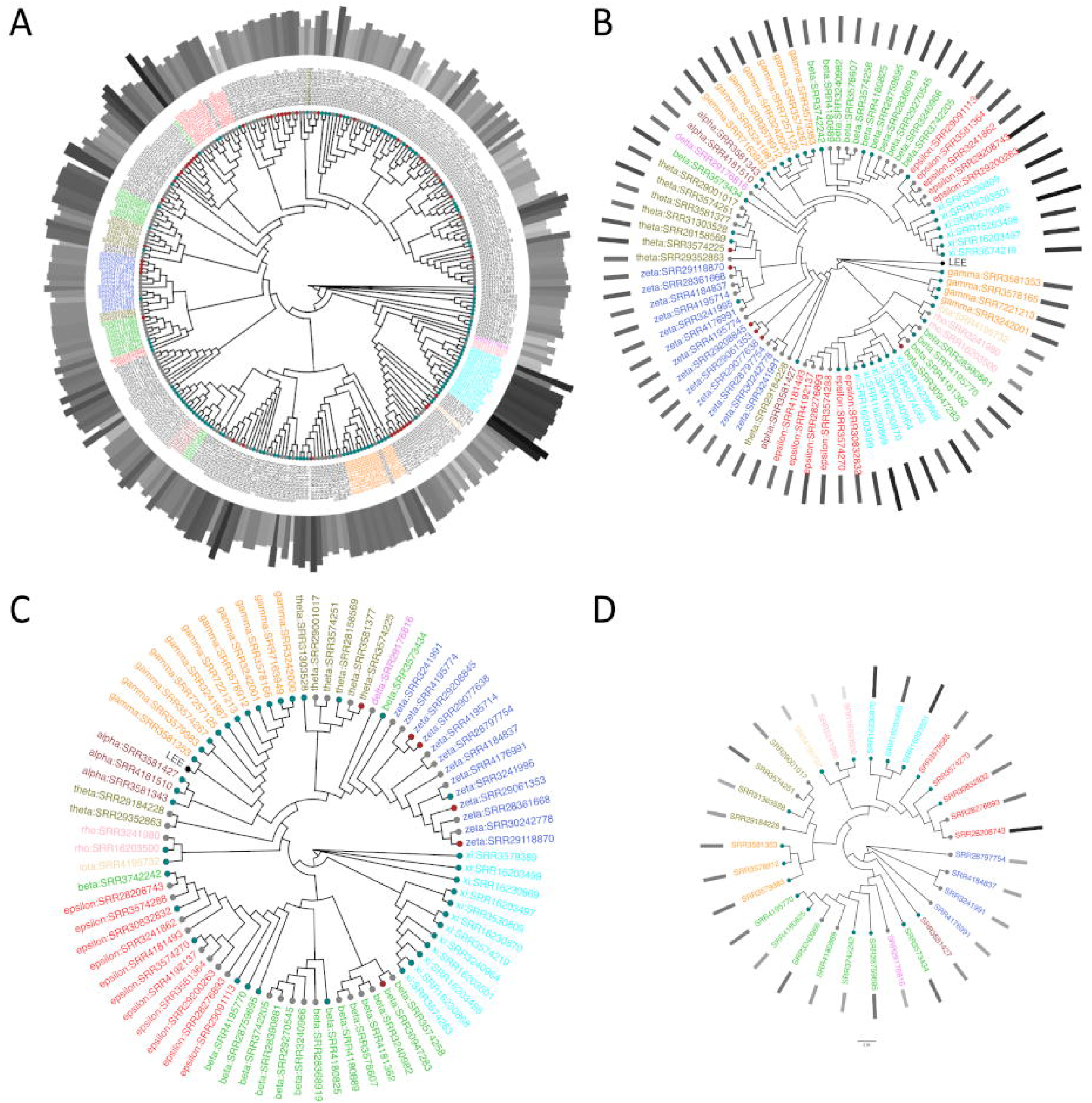
Phylogenetic tree for **A**. Core genome of 286 non-O157 STEC isolates, **B**. 85 *eae*+ LEE sequences. The LEE sequence from STEC O157 Sakai is labeled as ‘LEE’ in the phylogeny. **C.** Secretin C protein from the LEE of 85 *eae*+ isolates. Secretin C protein from the STEC O157 Sakai is labeled as ‘LEE’ in the phylogeny, and **D**. 31 unique *eae* sequences from *eae +*isolates (n=85). The leaf labels are colored according to the *eae* subtypes (eae negative:black, alpha:brown, beta:limegreen, gamma:orange, delta:magenta, rho:pink, iota:orange, delta:violet, epsilon:red, iota:wheat, theta:yellow, xi:cyan, zeta:royalblue), the bar length depicts the number of virulence factors present in the genome and the leaf node color is colored according to the Shiga Toxin subtype (*stx1*: grey, *stx2*: teal, *stx1* and *stx2*: brown).

The phylogeny obtained using the full-length LEE sequences from 85 *stx*+/*eae*+ isolates (Figure 4B) results in better clustering of the *eae* subtypes compared to the core genome phylogeny, confirming the fundamental role of genomic location. Further, it is interesting to note that the *eae* subtypes cluster even better in a phylogeny derived using only the LEE-located gene for the Type 3 secretion system Secretin (Figure 4C). This gene is essential to the function of T3SS, and its evolution is a good measure of that of the LEE island. Identifying the core genome of the LEE island using pangenome analysis (as previously applied to the whole bacterial genome), and constructing a phylogeny based on the set of core genes, produces similar results (Figure S3A). These phylogenies show that evolution of intimin subtypes largely follows evolution of the LEE island, except for some splitting of the beta and theta clades. Finally, a phylogeny was constructed from the 31 unique translated intimin sequences, reflecting only nonsynonymous variations (Figure 4D). This shows perfect separation of intimin subtypes, except for the inclusion of the single delta intimin sequence within the beta clade.

Next, the evolutionary mechanism of the intimin gene was investigated. While point mutations produce branches in a tree-like pattern, recombination events lead to networks with closed loops. Networks were generated using split decomposition analyses of the full length (Figure S3B), N terminus (1-700 amino acids, Figure S3C) and C-terminus (701-939 amino acids including the D2 and D3 domains, Figure S3D) of the intimin sequences, using SplitsTree^40^. These network patterns indicate horizontal gene transfer through recombination is more prominent in the N-terminus of the intimin sequence. Split decomposition analysis using an alternative division into N-terminus (1-550) and C-terminus (551-939 including domains D0, D1, D2 and D3) regions showed a similar trend, with the network using the N-terminus region more reticulate than the C-terminus region (Figure S4).

### 3.3 Structural analysis of exemplar virulence factors: Intimin-Tir

#### 3.3.1 Variations in Intimin

Allelic variation in *eae* was shown to be associated with STEC genotype, indicating a degree of clonality^53^. While there are no experimental structures of the full-length Intimin and Tir proteins, there are crystal or NMR structures of individual domains. Intimin is usually ∼939 amino acids long and comprises a flexible periplasmic N-terminal region (up to ∼188 amino acids), followed by the transmembrane beta-barrel domain (189-549) and the extracellular C-terminal region (550-939) (Figure 5A). The super-domain D2/D3 in the C-terminal region is responsible for binding the cognate receptor Tir and forming host attachment. We use residue numbering based on the UniProt reference sequences P19809 (Eae) and Q9KWH9 (Tir), which are represented by the structure PDB ID: 1F02^54^. This reference protein sequence for Intimin has 99.79% identity with the translated sequence of an alpha subtype *eae* annotated in Virulence Finder, although other reference alpha subtypes from Virulence Finder have lower identities down to 86% (illustrating variation within subtypes). The P19809 sequence also clusters with alpha subtype sequences within our set of isolates.

**Figure 5:**
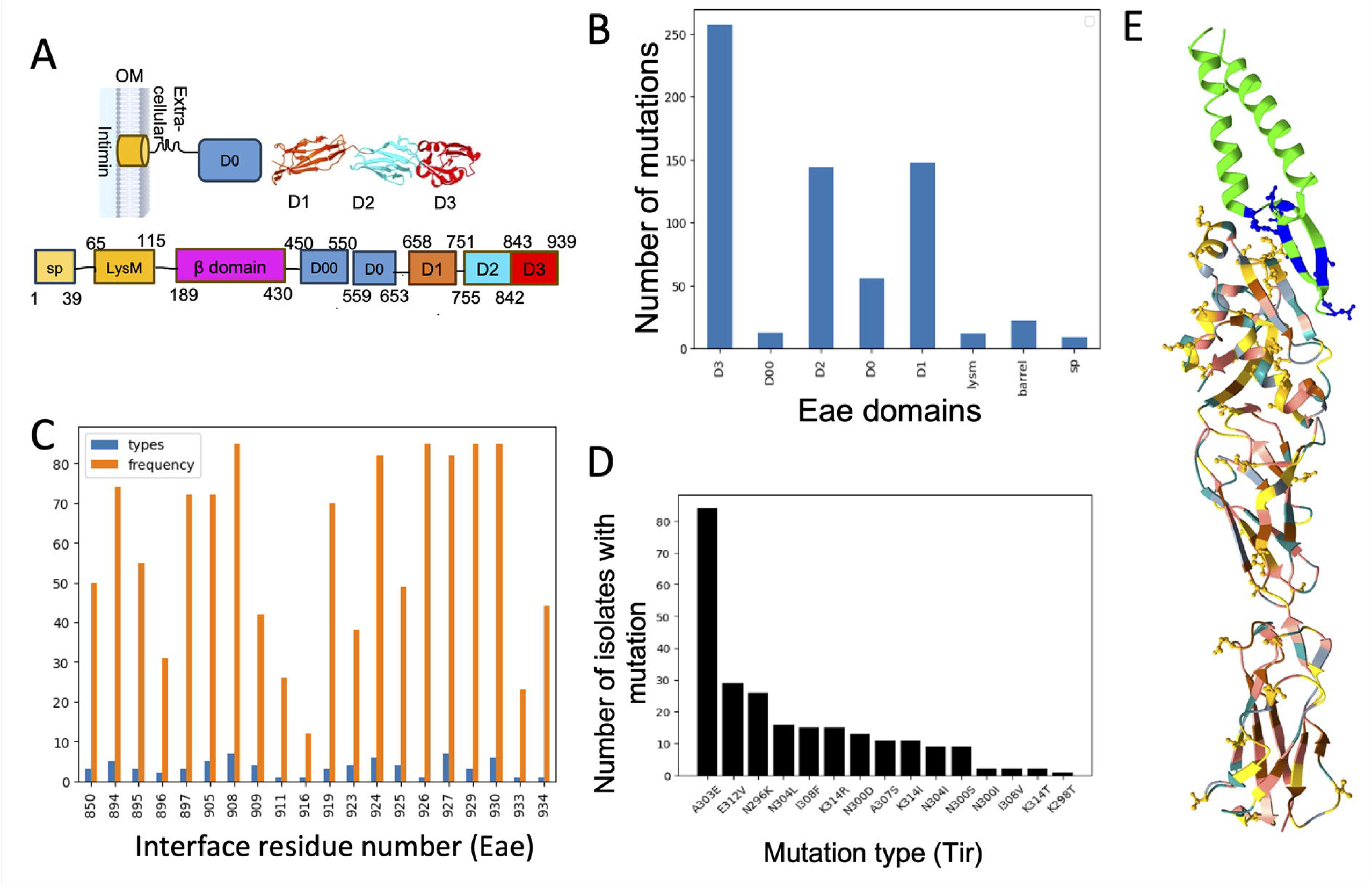
Variations in the intimin amino acid sequences. **A**. Schematic of Eae domains, domain boundaries are marked. **B**. Number of variations in different intimin domains. **C**. Number and types of variations of interface residues present in Eae’s D3. The number of Eae sequences with mutation is shown as orange bar and the height of blue bar denotes the types of variations observed for that particular interface residue. **D.** 15 types of variations in the beta hairpin region of Tir, which is known to interact with Eae. **E.** Structural mapping of variations observed in Eae (salmon) and the Eae-binding region of Tir (green). The variations in Eae are colored according to their frequency; 1< frequency <=20 (brown), 21< frequency <=40 (gray), 41< frequency <=60 (teal), 61< frequency <=80 (yellow) and frequency >80 (golden sticks).

The 85 intimin protein sequences from our *stx*+*/eae*+ isolates (comprising 31 unique sequences) were aligned to the UniProt sequence P19809 (subtype alpha), resulting in the identification of 706 point variations. These were mapped onto the domains using the following domain boundaries: 1-39, 65-115, 189-430, 450-550, 559-653, 658-751, 755-842, 843-939, corresponding to the SP (Signal Peptide), LysM, barrel, D00, D0, D1, D2 and D3 domains respectively (Figure 5A). Overall 661/706 variations lie within these domains, with the rest present in linker regions, and the majority of these (78%) lie in the exposed domains D1, D2 and D3 (Figure 5B).

Residues that contribute to the interface with Tir are different in our isolates to those observed in the P19809 reference sequence (PDB ID: 1F02), with a limited number of variant types permitted (Figure 5C, Table 2). Residue 926 is mutated to a valine in all 85 *eae*+ isolates, including those identified as alpha subtype, suggesting a global shift rather than variability. On the other hand, the neighbouring residue 927 shows 8 residue types (including the reference Lys) suggesting more variability at this site. Some of these variants involve a change in the amino acid type and side chain length such as (I897T, I897S, T895I), whereas others involve more subtle physicochemical changes such as I926V, K919R, Y934F, R850K. The variations at the interface are investigated in more detail below (section 3.3.4).

**Table 2:** Amino acid variations observed in the interface residues in Eae sequences from the dataset of *stx*+/*eae*+ isolates. Residues are ordered according to their observed variability, namely the number of distinct variations with respect to the reference sequence P19809.

| Reference residue (1F02) | Types of variation | Number of isolates with ref residue mutated | Mutated residue(s) |
| --- | --- | --- | --- |
| K908 | 7 | 85 | S, R, T, Q, N, I, P |
| K927 | 7 | 82 | G, D, H, T, E, S, N |
| N924 | 6 | 82 | P, T, I, A, Q, - |
| E930 | 6 | 85 | K, A, T, N, D, S |
| Q894 | 5 | 74 | K, L, R, E, N |
| Q905 | 5 | 72 | D, A, V, K, S |
| S909 | 4 | 42 | Q, G, E, K |
| N925 | 4 | 49 | G, S, D, R |
| L923 | 4 | 38 | V, R, H, I |
| I897 | 3 | 72 | T, S, N |
| K919 | 3 | 70 | T, R, A |
| T895 | 3 | 55 | S, A, I |
| R850 | 3 | 50 | K, Q, P |
| S929 | 3 | 85 | K, N, T |
| I896 | 2 | 31 | L, M |
| I926 | 1 | 85 | V |
| Y934 | 1 | 44 | F |
| V911 | 1 | 26 | W |
| A933 | 1 | 23 | V |
| D916 | 1 | 12 | N |

#### 3.3.2 Variations in Tir (translocated intimin receptor) protein sequence

The cognate receptor of intimin, Tir, is known to be rich in disordered regions, with flexible intracellular domains (*i.e.* N-tir and C-tir) that allow it to engage with a variety of host proteins^55^. Tir binds Intimin via its extracellular domain (residues 271-336, using the sequence numbering of PDB ID: 1F02) with much of the interaction mediated by a beta-hairpin (residues 296-315)^56^.

The extracellular domain of Tir was observed to be more conserved than the Tir binding region of Eae. There were 49 variant types of which 15 types are in the beta hairpin of Tir located at 9 distinct sites (Figure 5D). All of these variations in beta hairpin were present in less than 50% of the genomes except for the A303E which was observed in nearly all (84) of the eae+ sequences (Figure 5D).

#### 3.3.3. Structural mapping of Eae variations

706 variant types identified in our set of eae+ isolates (section 3.3.1), map to 320 residues. AlphaFold3 was used to model the full length structure of Eae (UniProt ID:P19809, PDBID: 1F02) and these 320 residues (in black sticks) were mapped onto the modelled structure (Figure S5A). The N-terminal region is modeled with low confidence as indicated by poor pLDDT scores (<50 or between 50-70). Additionally, higher PAE (predicted aligned error) scores (Figure S5B) suggests that AlphaFold3 is uncertain about the relative positioning of the domains (except for D1,D2 and D3 domains) in the modeled structure. The mapping of observed mutations suggests most of the variations are in the extracellular regions of Eae. This poor quality, low confidence structure is only used to map the sequence variations onto the structure and this low quality structure is not used for drawing any functional or structural implication of these variations.

The reference crystal structure (PDB ID:1F02) corresponds to domains D1, D2 and D3 (residues 658-939) of Eae along with the Eae binding region of Tir (residues 271-336), and we mapped the point variations observed in these regions onto this structure. Figure 5E shows the crystal structure, with Eae coloured according to the frequency of mutation with respect to the sequence of the crystal structure (P19809). The most frequently mutated residues (observed in > 80 isolates) are shown in gold ball-and-stick representation. As discussed above, these mutations consist of one or more mutation types. Of these high frequency mutations, 55% were observed in the D3 domain (Tir binding domain). The variations in the interface residues of Tir (residues 296-315) are shown as blue sticks (Figure 5E).

#### 3.3.4 Structural modelling of intimin subtypes

In order to investigate the functional consequences of sequence variation, the structures of Intimin-Tir complexes were modelled using AlphaFold3^27^. As noted previously (section 3.2.2) there are ten *eae* genetic subtypes represented in our set of *stx*+/*eae*+ isolates. Therefore, ten structural models were generated corresponding to these *eae* subtypes in complex with their cognate Tir partner. The quality of the modelled complexes is indicated by the ipTM (interface Predicted TM) score, which measures the accuracy of the predicted positions of the subunits in the modelled complex. ipTM scores greater than 0.8 are predictions with high confidence and predictions with ipTM <0.6 are failed predictions^27^. All the modelled complexes except rho and theta have high confidence predictions for the relative subunit positions (Table 3). Therefore for rho and theta subtypes, homology modelling (Modeller v 9.10) was used to generate the Eae-Tir structural complex using PDB structure 1F02 as a template.

**Table 3:** Confidence scores for the Eae-Tir modelled complexes. All the complexes except for rho and theta were modelled using AlphaFold3 (AF3). For the subtypes rho and theta, homology modelling was used to model the complex using PDB structure 1F02 as a template. *Interface scores were obtained for AF3 models for all subtypes except rho and theta, for which it is calculated on the homology models generated using Modeller.

| Intimin Subtype | ipTM score (AF3 models) | Interface score |
| --- | --- | --- |
| gamma | 0.88 | 0.98 |
| alpha | 0.83 | 0.98 |
| beta | 0.82 | 0.96 |
| delta | 0.79 | 0.98 |
| epsilon | 0.81 | 0.98 |
| iota | 0.81 | 0.99 |
| rho | 0.34 | 0.99* |
| theta | 0.17 | 0.99* |
| xi | 0.83 | 0.98 |
| zeta | 0.83 | 0.99 |

Furthermore, the quality of the Eae-Tir interface of these modelled complexes was assessed using an in-house deep learning based method (DPI-score, Bhujel *et al.*, manuscript under preparation). All the modelled complexes were predicted to have a high quality score for the protein-protein interface (Table 3).

Upon validation of the quality of modelled complexes, the effects of genomic variations that were observed at the interface between Intimin and Tir were investigated. The variants present at the interface between Intimin and its cognate Tir were assessed in order to elucidate the gain of interactions required to preserve the receptor binding and maintain function. Some of these co-occurring variations (including two, E312 and N300, of three Eae binding hotspots as listed in Ross and Miller, 2007^56^) and their impacts are highlighted here.

E312V(Tir):

E312(Tir) is a known crucial interface residue (PDB ID: 1F02, Figure 6A) and is identified as one of the intimin binding hotspot residues^56^. In the reference crystal structure, it is observed to interact with K927(Eae) and K298(Tir). In the zeta, alpha and gamma intimin subtypes in our set of non-O157 *eae*+ isolates, their cognate Tir protein sequences were observed to have the mutation E312V (Figures 6B, 6C and 6D respectively). E312V results in the loss of these interactions due to the non-polar side chain of valine.

**Figure 6:**
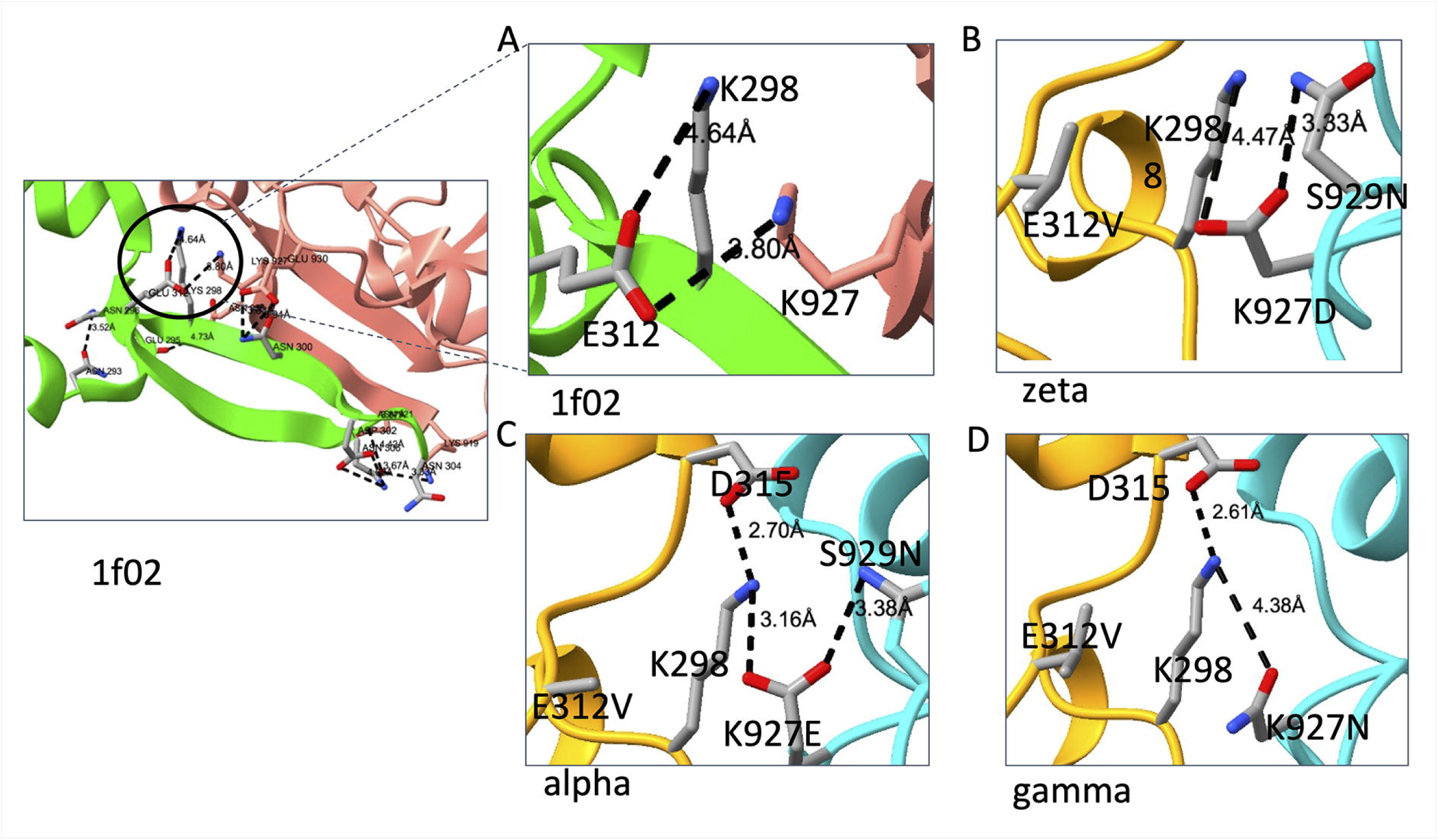
Structural inspection of interactions between Eae-Tir (Tir:E312 and Eae: K927 and K 928). **A**. As in the crystal structure, 1F02, the residue E312(tir) sits at the interface with Eae and interacts with K298 and K927. **B**. In the zeta subtype, E312V results in the loss of these interactions but simultaneously the gain of interactions is observed due to the other variations in *eae* namely K927D and S929N to preserve the interactions at the interface. **C.** In the alpha subtype,the co-occurring variations K927E and S929N in Eae were observed to gain interactions at the interface. **D**. In the gamma subtype, E312V, there were no notable gain in interactions.

However, to counter the effects of this loss of interactions, K927D, K927E and K927N were observed in the zeta, alpha and gamma *eae* subtypes respectively (Figure 6B-D). The neighboring residue, K298(Tir), is a conserved residue and plays a role in preserving the Eae-Tir interactions in the *eae* subtypes. Additionally, S929(Eae) was observed to be mutated to S929N(Eae) in these three subtypes, which help to preserve the interactions with the conserved residue K298. However, out of the three variants, gamma has the least number of compensatory interactions that involve K298 (Figure 6D), although there may be compensatory changes at other interaction sites.

##### N300(Tir)

N300(Tir) is another Eae intimin binding hotspot^56^ and may form a hydrogen-bond with E930 in the crystal structure (PDB ID: 1F02, Figure 7A). N300(Tir) was observed to be mutated to D, I or S in the isolates having *eae* subtypes zeta, rho and theta respectively. Upon structural inspection, N300D appears to form a salt bridge with E930K in the zeta subtypes (Figure 7B). This pair of co-occurring mutations shows a reversal of residue type, with the glutamic acid of 930(Eae) being replaced by the aspartic acid of 300(Tir), and illustrates the compensation required to maintain function, that is Eae-Tir binding.

**Figure 7:**
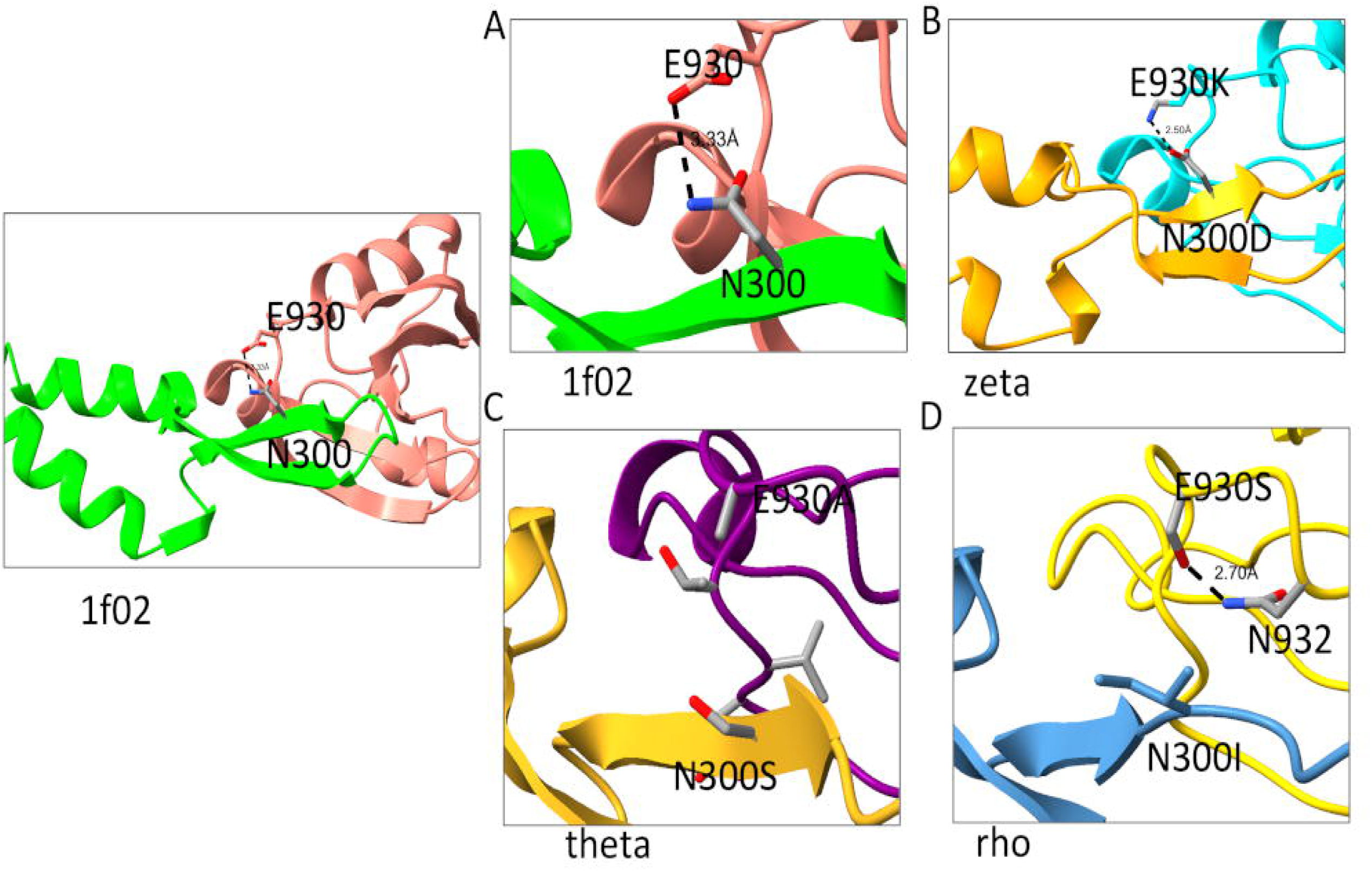
Structural inspection of interactions between Eae-Tir (Tir: N300 and Eae: E930). **A**. N300 is seen to be in proximity to E930 and they may be involved in H-bond. **B.** In zeta subtype, this interaction is seen to be preserved by the formation of salt bridge between the residues N300D and E930K. **C**. In theta subtype, N300S and E930A are observed however no specific gain of interaction is observed upon the mutation. **D**. In the rho subtype, N300I is observed along with E930S. E930S may be forming a H-bond with N932 to maintain the structural stability of the Eae.

The structural models for rho and theta produced using AlphaFold 3 predicted a different interaction site between Eae and Tir than the one observed in the crystal structure (1F02, Figures S6A and S6B). To try to obtain the expected quaternary structure, we modelled the complex with AlphaFold 3 including only the D3 region of the Eae subtype sequence together with the Tir extracellular domain. However, superposition of the predicted complex structure with the crystal structure showed many changes in the secondary structure of Eae which are probably artefacts due to the loss of D2 (Figure S6C and S6D). Therefore, it was necessary to resort to the traditional approach of homology modelling using Modeller^41^ (v10.4) and the crystal structure as template.

These homology models were used to examine the variations in the rho and theta subtypes. There did not appear to be any major impacts of N300S in the theta subtype on the interface between Eae and Tir (Figure 7C), despite the loss of a specific interaction with Eae. E930A is the co-occurring mutation observed in its Eae sequence. For the rho model, no gains of interaction were observed for N300I(Tir). However, the co-occurring mutation E930S(Eae) may form a H-bond with N932 (Eae) to counter the effect of variations and preserve the structure and function (Figure 7D).

## 4. Discussion

In this study, a dataset comprising 286 non-O157 STEC genomes sourced from EnteroBase was curated and analysed to determine whether the virulence factor complement coupled with allelic variation could be used to infer their role in pathogenicity. We used the premise that genetic variation that results in structural changes impacts protein function and potentially their contribution to virulence. A selection of isolates was made to represent a diverse range of serotypes, with highly prevalent and well-characterised serotypes such as O26 excluded to minimise bias. The presence of the *stx* toxin gene was used to define these isolates as STEC. The genomic diversity of the set of non-O157 STEC isolates is illustrated by the fact that the genomes share only 3119 core genes out of a pangenome of ∼36k genes.

The study took a two-step approach, to first investigate the individual and co-occurrence of genetic factors that are thought to contribute to disease, defined as Virulence Factors^24,25^. Secondly, since the majority of virulence factors exist in different allelic variants, the potential functional impact of variation was investigated, focusing on the well characterised STEC virulence factor Intimin, Eae and its cognate partner, Tir. Variations were considered both at the subtype level and at the level of individual point mutations. This serves as an example for other protein interactions of virulence factor variants, and potential impacts on overall pathogenicity of non-O157 STEC isolates.

The non-O157 STEC are characterised by their genetic diversity. Using serotype alone as a distinguishing feature identified 111 variants in our dataset. Like other members of the *E. coli* species, it is evident that the STEC sub-species comprise mosaic genomes, with multiple instances of recombination, genomic rearrangements and horizontally acquired elements, making classification challenging. The decision tree from the JEMRA risk matrix is based on the *stx* gene subtype and the presence of the intimin gene, *eae*, and the transcriptional activator of aggregative adherence fimbria expression, *aggR*. However, the intimin gene is not ubiquitous in clinical samples and was only present in 85 / 286 (30 %) *stx*+ non-O157 STEC genomes investigated. Furthermore, *aggR* is not widespread in EHEC and does not occur at all in our *stx+* set.

Some serotypes of STEC that lack initimin are notable as they still cause severe disease^57^. STEC O91 is reported to be one of the most common *eae*- non-O157 serotypes associated with human disease^58^. O91 and O113 have been reported as prevalent serotypes (top 9) in England in 2019^59^, and are represented in our dataset. STEC O55 outbreaks have also been reported in England^41^, represented by nine *eae*+ isolates and two *eae*- isolates in the dataset .

Besides *stx*, and *eae*, additional virulence factors are associated with pathogenicity of non-O157 STECs, and can contribute to severe disease phenotypes such as bloody diarrhoea and HUS. Genes that help with adherence, colonisation, production and secretion of toxins are known to be associated with causing severe disease. Within our dataset, there was a large degree of variation in the complement of virulence factors, with 130 different types of virulence factors across the set. Some factors are present in most isolates (e.g. *nlpI, terC, yeh, csgA*), while others are only represented in single isolates, such as *csm* and *fot*. The virulence factors that occurred frequently are known to be abundant in *E. coli* genomes^60^. They can be considered as factors that facilitate host colonisation or adhesion to a surface, and in doing so contribute to pathogenicity and lead to disease.

The co-occurrence of some pairs of virulence factors was evident, while other pairs were apparently mutually exclusive in the set of isolates. Several different sets of genes co-occurred, with the largest group encompassing a range of functions associated with virulence, including: *ireA*, a siderophore associated with pathology in UTI^61^ ; capsular polysaccharide production genes, *kps*^62^; microcin genes, *mchC* and *F,* associated with virulence in the STEC O91 serogroup^63^; the *Shigella* enterotoxin gene *sen,* also associated with bacteraemic *E. coli* isolates^64^; *shiA,* responsible for immunomodulatory functions in *Shigella* species^65^, with homologs apparently with similar functions in uropathogenic *E. coli*^66^; the subtilis cytotoxin gene *subA*, associated with STEC^67^ including isolates from cattle^68^; and the associated *tia* gene, shown to be involved in cellular invasion of some STEC strains^69^. Notably, the latter two virulence factors are encoded on the subtilase-encoding pathogenicity island (SE-PAI) and are mutually exclusive with the locus for enterocyte effacement^70^, presumably because of the role in invasion as opposed to A/E lesion formation.

Co-occurrence of virulence factors with the initimin gene, *eae*, was quantified to obtain a clearer indication of potentially interacting networks based on this key virulence factor. 52 % and 41 % of the *eae*+ isolates were *stx2* and *stx1*, respectively, with only 7 % occurring with both *stx*1+ and *stx*2+. As expected, some virulence factors are co-located with intimin on the LEE island e.g. *tir*, *espA, espB, espF*, although not all. This is in-line with the concept of a network of T3SS effectors, which exhibit co-dependency and context-dependent functionality^71^. Virulence factors that exhibited strong co-occurrence with *eae* include the non-LEE encoded T3SS effectors *nleA/B/C* and *cif,* and the plasmid-encoded genes *esp, tox* and *katP*.

Several of the genes identified in co-occurrence networks of virulence factors were first identified on the pO157 plasmid, which was genetically characterised in serotype O157:H7, strain Sakai^72^. It encodes genes for a type 2 secretion system and the alpha-hemolysin gene cluster, *hyl*^73^,as well as its associated protease, *katP*. *katP* has been detected in multiple non-O157 isolates on other plasmids, e.g. in O111^74^ and O26^75^, with sub-type a in particular associated with clinical disease^76^. It is normally linked with other plasmid encoded virulence factors: the heamolysin gene cluster (*hyl*), a lipid A modification system (*ecf* operon) and the serine protease (*espP*), although they do not always co-occur^74^. The *katP* gene is normally associated with LEE-encoding STEC strains, and not in non-LEE strains, presumably down to genetic incompatibility^77^. The KatP protease is responsible for cleavage and inactivation of Hyl heamolysin. STEC isolates frequently encode multiple hemolysins, of which four have been reported: alpha-hemolysin (HylA), enterohemolysin (ExhA) also encoded on a plasmid or phage (e-HylA) and silent hemolysin / cytolysin A (SheA)^78^. Both HylA and ExhA were detected in the STEC genome dataset. Hemolysins are encoded in other pathotypes of *E. coli*, although there has been shown to be a strong association between both ExhA and HylA and presence of *stx* genes and to some extent, with presence of *eae*^78^.

Using reference sequences taken from the Virulence Finder database, ten *eae* genetic subtypes were identified across the dataset of 85 isolates. Multiple subtypes of the *eae* gene have been reported previously, and correlated with host specificity and tissue tropism^79^. The *eae* gene was initially sequenced from EPEC lineage 1 strain E2348/69, forming the reference sequence for α-intimin, while EPEC lineage 2 strains encode subtype β-intimin. Further subtypes are defined for EHEC, such as γ-intimin from O157:H7 and O55:H7 strains^80^. Divergent intimins appear to be encoded by mosaic alleles composed of DNA segments with different evolutionary histories^81^.

Variation in intimin subtypes has long been recognised^81^, with at least 18 different genetic variants identified in a single non-O157 STEC genome set^79^. The *eae* variants identified here showed significant identity amongst themselves, with the epsilon and xi subtypes sharing ∼97% identity. Recombination events have been implicated in the evolution of *eae*, and split decomposition analysis on this dataset suggested a reticulate network indicative of recombination. The genomic variations (non-synonymous) in intimin subtypes were inspected using the sequence of *eae*-α as reference. ∼700 variations were observed in all intimin protein sequences and 78% of these lie in the extracellular region of intimin (domains D1, D2 and D3), which interacts with the receptor Tir. However, the extracellular region of Tir was observed to be more conserved than the extracellular region of intimin.

Protein structural modelling enabled a better understanding of the functional implications of the genomic variations. AlphaFold3^27^ was used to generate the three-dimensional structural models of the intimin subtypes with their cognate Tir receptor, validated with an in-house machine learning based method. With AlphaFold3, monomeric protein structures are modelled with high accuracy although protein-protein complexes modelling is still a developing analytical tool^82^. As such, three-dimensional structure modelled complexes need to be validated using confidence metrics such as pLDDT (per atom confidence score), ipTM (confidence in relative placement of subunits in a modelled complex). Although the structures are models and the atom positions are approximate, it nonetheless still provides a good estimate of interactions at the interface.

Mapping of Eae variations onto structural models shows that most occur in the extra-cellular domains, and of these most are in the interaction region with the cognate partner Tir. Mutations in the extra-cellular region may be associated with immune escape, but nevertheless binding with Tir must be maintained in order to retain effective pedestal formation. For some specific examples, we have shown that variations preserve the functional interaction between the two protein partners through a compensatory mechanism, wherein loss of some interactions at the interface is compensated by the formation of new interfacial contacts. Understanding the importance of specific interactions involved in receptor binding will help improve the interpretation of STEC genomic variants and may aid the development of interventions.

Through this work, we would like to emphasise the importance of integrating genomic analysis with protein structural analysis which provides insights into mechanism of function, in this case pathogenicity of eae+ STEC isolates. The examples of co-occurring variations at protein-protein interfaces underscore the importance of integrating genomic data with structural biology to deepen our understanding of protein function.

## Author Contributions

Sony Malhotra- Conceptualization, Formal analysis, Funding acquisition, Methodology, Writing-original draft, Writing- review and editing

Ashley Ward- Data curation

Lauren Giles- Data curation, Writing- review and editing

Tom Gerrard- Formal Analysis

Martyn Winn- Conceptualization, Formal analysis, Funding acquisition, Methodology, Writing-original draft, Writing- review and editing

Nicola Holden- Conceptualization, Data curation, Funding acquisition, Methodology, Project Administration, Writing-original draft, Writing- review and editing

## Supplementary Material

**Figure S1:** Co-occurrence of selected gene groups. Gene group ‘*aal*’ includes 4 genes, ‘*fae*’ includes 6 genes, and ‘*mch*’ includes 2 genes, giving a total of 32 genes.

**Figure S2:** Pairwise percent identity between the protein sequences of 10 intimin subtypes plotted as a heatmap.

**Figure S3: A**. The phylogeny obtained using the aligned core genome sequences of the LEE obtained from the *eae+* isolates (n = 85). Split decomposition network tree using **B**. full length intimin sequences, **C**. N-terminus region (1-700) and **D**. C-terminus region (701-939) of the intimin sequences.

**Figure S4:** Split decomposition network tree using **A**. N-terminus region (1-550) and **B**. C-terminus region of the intimin sequences.

**Figure S5:** Full length structural model of Intimin. **A**. Full length structural model for the UniProt sequence P19809 (Intimin) obtained using AlphaFold3. The sequence domains are colored: sp(yellow), LysM(orange), β-barrel(magenta), D0 and D00(blue), D1 (orange red), D2(cyan) and D3 (red). The point mutation sites are shown as black sticks. **B**. PAE (predicted aligned error) plot of the structural model obtained using AlphaFold3.

**Figure S6:** Different structure prediction approaches for Eae-Tir sequences. The crystal structure (PDB ID: 1F02) in all the panels is shown in green (Tir) and salmon (Eae). **A**. Structural model of the translated rho Eae-Tir (blue) complex modelled using AlphaFold3 is superposed over the crystal structure. The interface modelled for the rho subtype is different to the one observed in crystal structure. **B**. Structural model of only D3 region of rho subtype with its cognate Tir built using AlphaFold3. The modelled complex is in blue. **C.** The structural model of the theta Eae-Tir (yellow) complex modelled using AlphaFold3 is superposed over the crystal structure. The interface modelled for the theta subtype is different to the one observed in crystal structure. **D**. Structural model of only D3 region (yellow) of theta subtype with its cognate Tir built using AlphaFold3.

**Table S1:** Distribution of non-O157 STEC serotypes in the human isolates in the UK obtained from Enterobase.

**Table S2:** Accession IDs (Enterobase) of the non-O157 STEC isolates used in the study.

**Table S3:** Virulence factor genes (using 3 letter nomenclature) in non-O157 STEC isolate genomes (n=286). Their presence in intimin positive isolates, enrichment factor and Jaccard similarity is also listed.

**Table S4:** Virulence factor genes (using 4 letter nomenclature) in non-O157 STEC isolate genomes (n=286). Their presence in intimin positive isolates, enrichment factor and Jaccard similarity is also listed.

