## Supplementary figures and images for "Stitching genomics data to protein structures: Virulence factors in non-O157 Shiga toxin-producing *Escherichia coli*"

### Supplementary Figure3

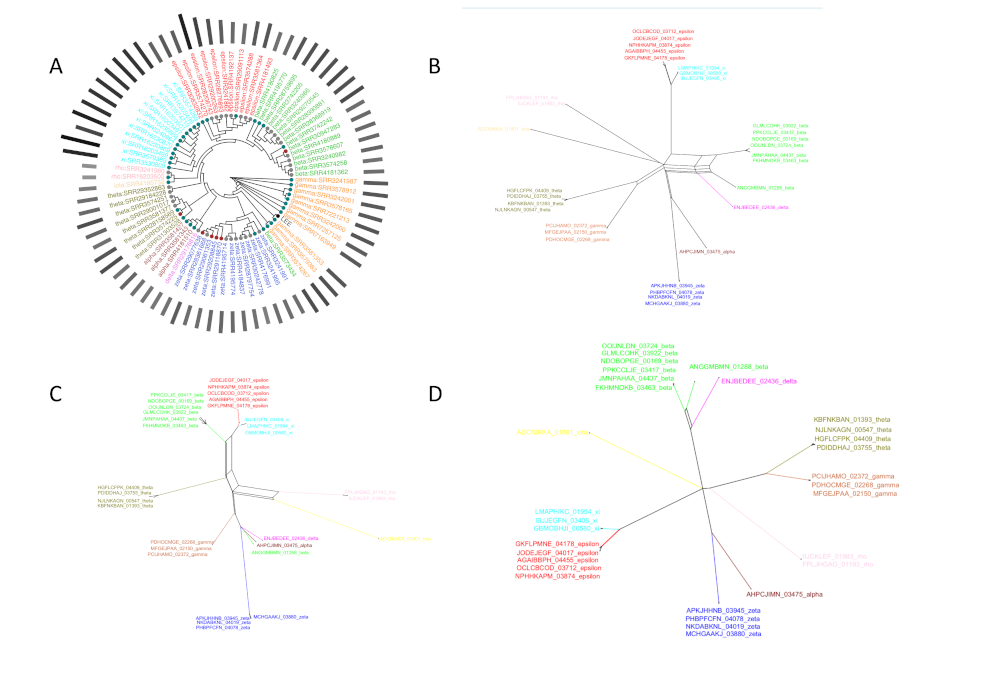

### Supplementary Figure 1

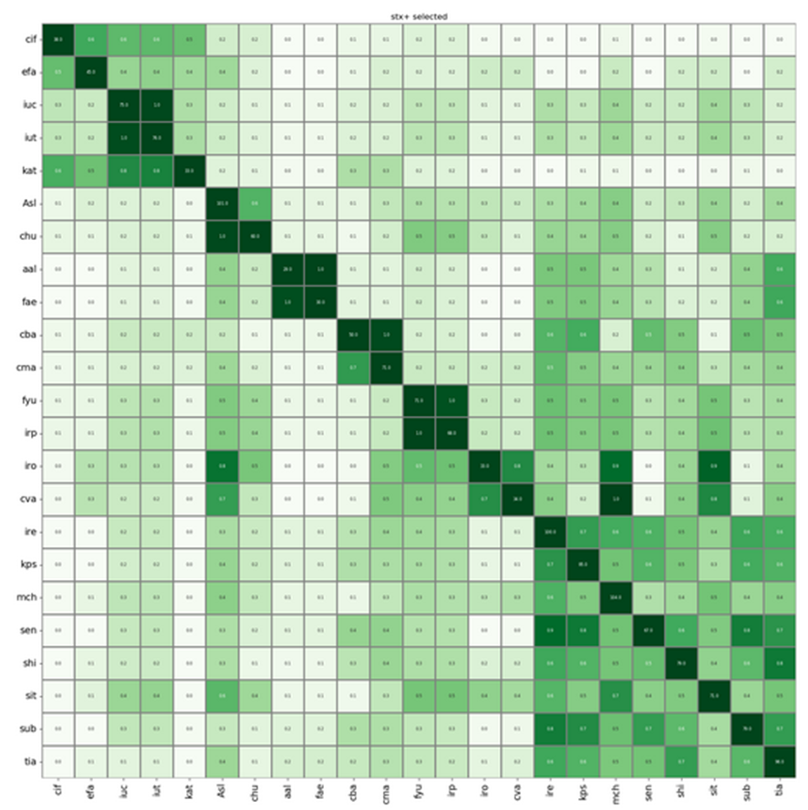

### Supplementary Figure 2

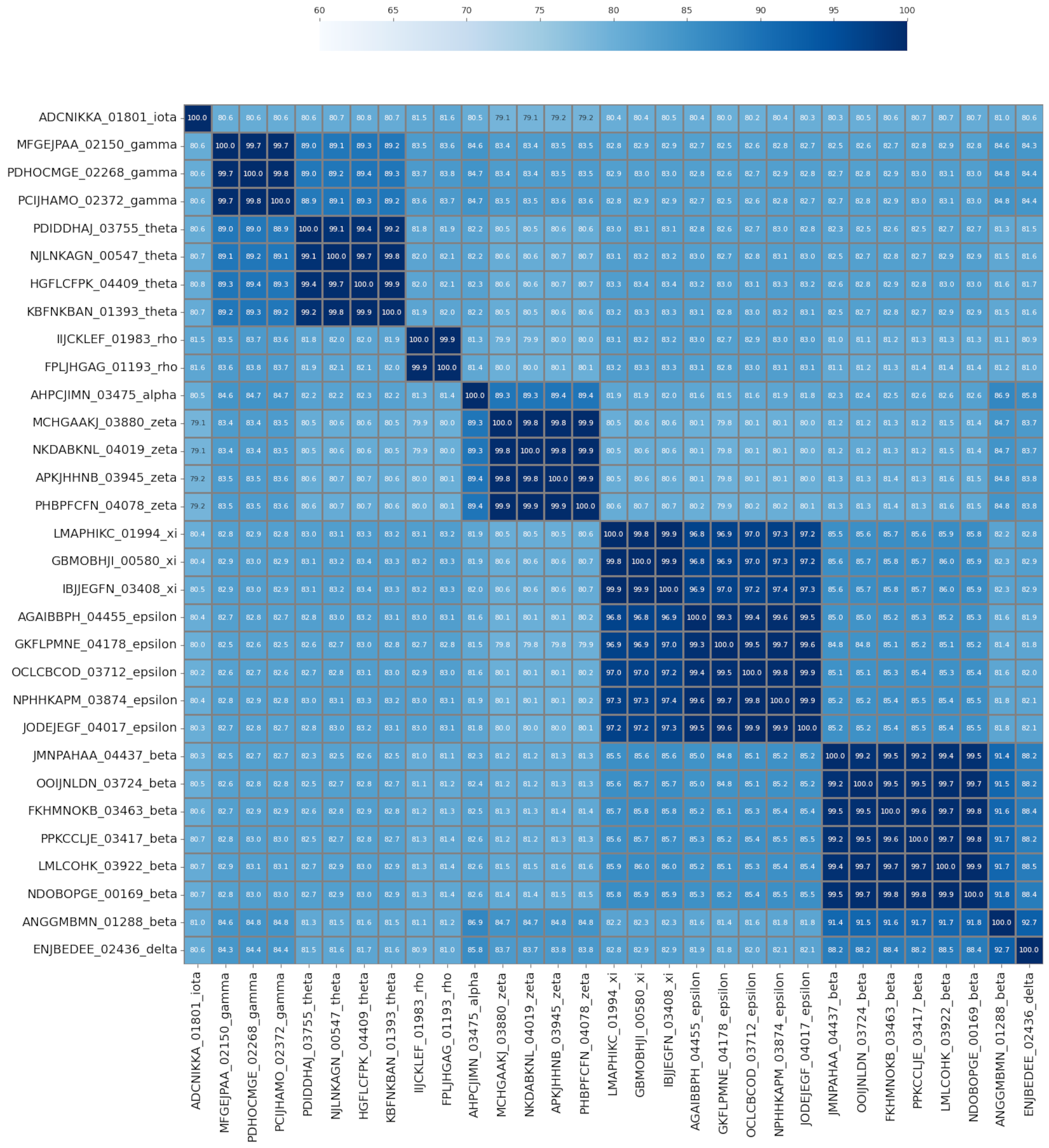

### Supplementary Figure 4

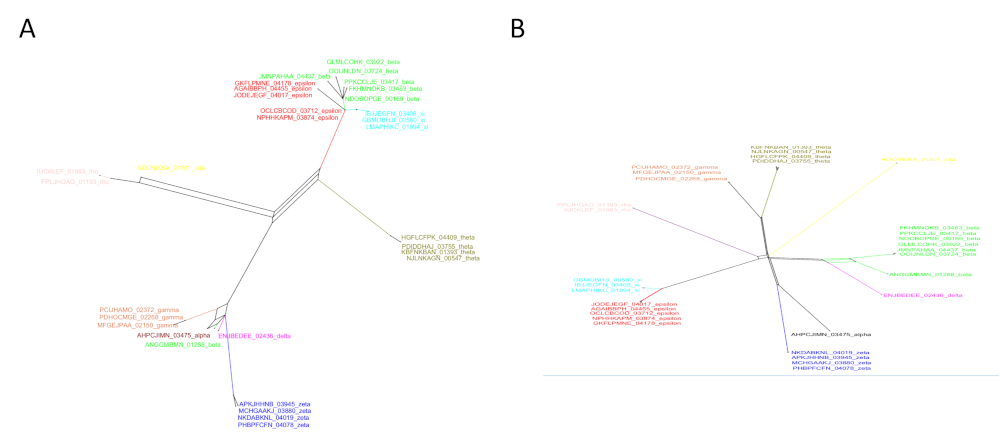

### Supplementary Figure 5

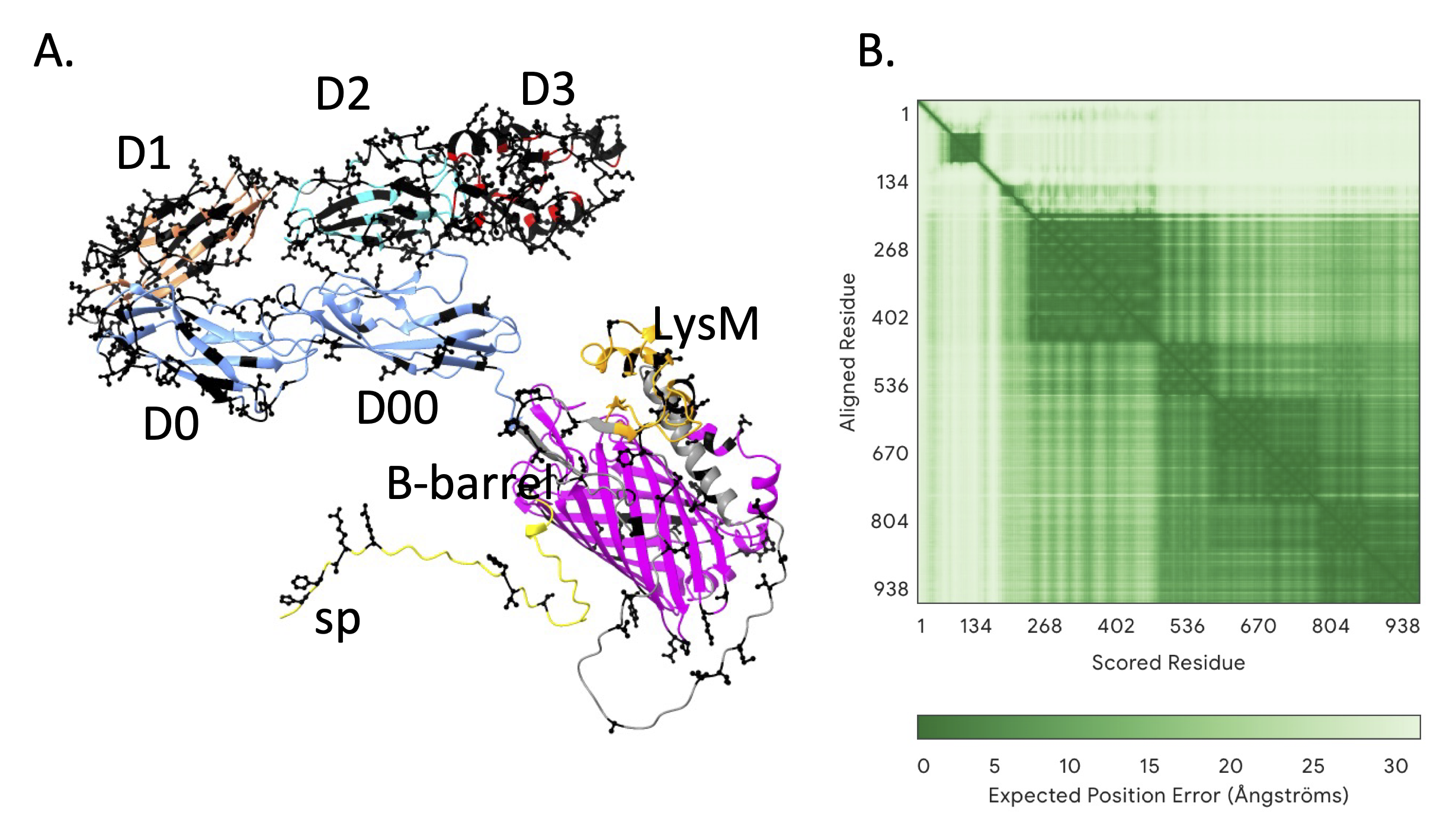

### Supplementary Figure 6

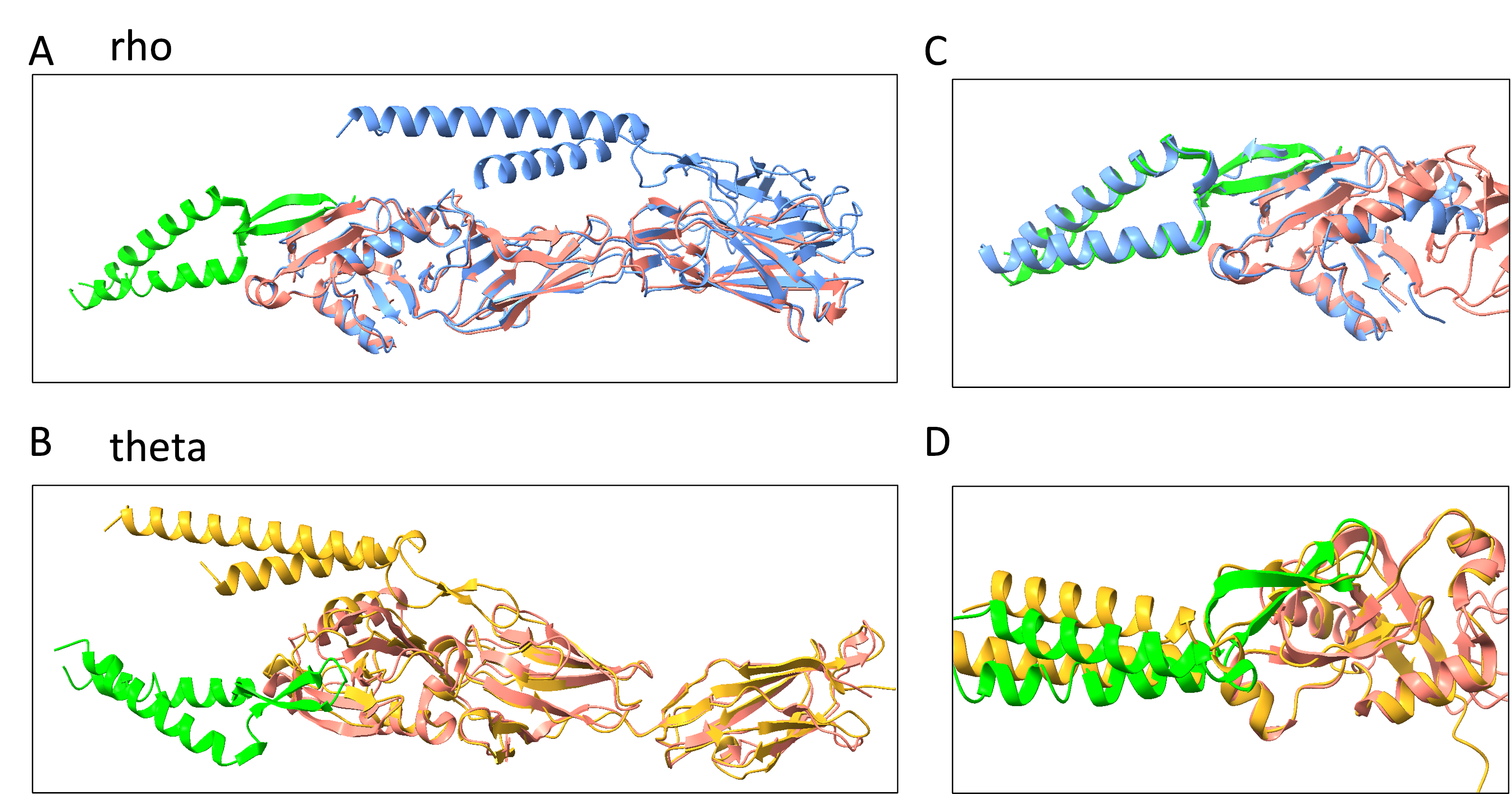
