## Supplementary Tables 1 and 2 for "Stitching genomics data to protein structures: Virulence factors in non-O157 Shiga toxin-producing *Escherichia coli*"

Table S1: Distribution of non-O157 serotypes in the human isolates in the UK obtained from Enterobase.

| Serotype | Count |
| --- | --- |
| O26:H11 | 250 |
| O145:H28 | 103 |
| O91:H14 | 68 |
| O146:H21 | 62 |
| O26 | 55 |
| O55:H7 | 49 |
| O25:H4 | 43 |
| O128Ab:H2 | 39 |

Table S2: Accession ids (Enterobase) of the non-O157 STEC isolates used in our study.

| SRR3242000 | SRR4184704 | SRR29176810 | SRR3578161 | SRR30664734 |
| --- | --- | --- | --- | --- |
| SRR3241995 | SRR4195788 | SRR29184250 | SRR3742205 | SRR30813998 |
| SRR3241993 | SRR4195770 | SRR29184200 | SRR4181529 | SRR30813976 |
| SRR3241992 | SRR4195732 | SRR29200070 | SRR4181493 | SRR30991897 |
| SRR3241989 | SRR4195719 | SRR29208845 | SRR4181441 | SRR31372593 |
| SRR3241987 | SRR4195714 | SRR29208603 | SRR4181362 | SRR31372591 |
| SRR3241983 | SRR4195622 | SRR29226799 | SRR4180897 | SRR31386880 |
| SRR3241857 | SRR4195473 | SRR29226023 | SRR4180889 | |
| SRR3241854 | SRR4192137 | SRR29252351 | SRR4179996 | |
| SRR3241848 | SRR7163949 | SRR29252336 | SRR4178298 | |
| SRR3241846 | SRR7221213 | SRR29241024 | SRR4176991 | |
| SRR3241834 | SRR7257125 | SRR29259443 | SRR4184858 | |
| SRR3241098 | SRR7588703 | SRR29259290 | SRR4184837 | |
| SRR3240980 | SRR16203497 | SRR29301502 | SRR4195774 | |
| SRR3240971 | SRR16203501 | SRR29648597 | SRR4195743 | |
| SRR3240968 | SRR16203498 | SRR29707989 | SRR4195476 | |
| SRR3240964 | SRR16203499 | SRR29707969 | SRR4195471 | |
| SRR3240961 | SRR16203500 | SRR29790549 | SRR4193853 | |
| SRR3531464 | SRR16230868 | SRR29908410 | SRR4192114 | |
| SRR3530809 | SRR16230869 | SRR30118407 | SRR28158799 | |
| SRR3530586 | SRR16230870 | SRR30104557 | SRR28158790 | |
| SRR3574336 | SRR27036283 | SRR30359106 | SRR28158768 | |
| SRR3574324 | SRR28158839 | SRR30359032 | SRR28220660 | |
| SRR3574270 | SRR28158787 | SRR30813946 | SRR28220655 | |
| SRR3574267 | SRR28158569 | SRR30832832 | SRR28208743 | |
| SRR3574264 | SRR28220739 | SRR30832784 | SRR28172130 | |
| SRR3574263 | SRR28220643 | SRR30947283 | SRR28232618 | |
| SRR3574242 | SRR28208692 | SRR31034849 | SRR28276893 | |
| SRR3574228 | SRR28200684 | SRR31047827 | SRR28284973 | |
| SRR3574225 | SRR28200608 | SRR31203340 | SRR28361675 | |
| SRR3574219 | SRR28172319 | SRR31294318 | SRR28361673 | |
| SRR3574026 | SRR28172302 | SRR31303528 | SRR28361668 | |
| SRR3573434 | SRR28172158 | SRR31372567 | SRR28349722 | |
| SRR3581502 | SRR28232614 | SRR3241999 | SRR28364854 | |
| SRR3581443 | SRR28361616 | SRR3241994 | SRR28368919 | |
| SRR3581433 | SRR28361614 | SRR3241991 | SRR28382969 | |
| SRR3581427 | SRR28382941 | SRR3241981 | SRR28382848 | |
| SRR3581426 | SRR28390887 | SRR3241980 | SRR28390900 | |
| SRR3581400 | SRR28402139 | SRR3241864 | SRR28390881 | |
| SRR3581360 | SRR28428450 | SRR3241862 | SRR28402113 | |
| SRR3581358 | SRR28428357 | SRR3241860 | SRR28621924 | |
| SRR3581353 | SRR28503468 | SRR3240982 | SRR28797754 | |
| SRR3581348 | SRR28511710 | SRR3240966 | SRR28825009 | |
| SRR3581343 | SRR28646855 | SRR3240965 | SRR28891380 | |
| SRR3581339 | SRR28759695 | SRR3574302 | SRR29001017 | |
| SRR3581324 | SRR28759640 | SRR3574288 | SRR29070903 | |
| SRR3581320 | SRR28759633 | SRR3574285 | SRR29070875 | |
| SRR3579390 | SRR28825084 | SRR3574258 | SRR29061320 | |
| SRR3579389 | SRR28894213 | SRR3574252 | SRR29080100 | |
| SRR3579383 | SRR29004345 | SRR3574251 | SRR29077767 | |
| SRR3578955 | SRR29046134 | SRR3574248 | SRR29091383 | |
| SRR3578915 | SRR29074272 | SRR3581423 | SRR29176898 | |
| SRR3578912 | SRR29074263 | SRR3581389 | SRR29176816 | |
| SRR3578798 | SRR29061392 | SRR3581382 | SRR29184228 | |
| SRR3578636 | SRR29061353 | SRR3581377 | SRR29200263 | |
| SRR3578585 | SRR29061246 | SRR3581364 | SRR29208598 | |
| SRR3578294 | SRR29080102 | SRR3579402 | SRR29225483 | |
| SRR3578165 | SRR29080067 | SRR3579387 | SRR29252521 | |
| SRR3742268 | SRR29077671 | SRR3578976 | SRR29252407 | |
| SRR3742242 | SRR29077638 | SRR3578919 | SRR29252335 | |
| SRR3742209 | SRR29091312 | SRR3578909 | SRR29241231 | |
| SRR4181510 | SRR29091113 | SRR3578784 | SRR29259424 | |
| SRR4180890 | SRR29091107 | SRR3578659 | SRR29270545 | |
| SRR4180825 | SRR29119977 | SRR3578635 | SRR29352863 | |
| SRR4180191 | SRR29118870 | SRR3578631 | SRR29672707 | |
| SRR4179920 | SRR29118867 | SRR3578607 | SRR30150752 | |
| SRR4192078 | SRR29154876 | SRR3578592 | SRR30242778 | |
| SRR4184861 | SRR29154844 | SRR3578167 | SRR30515325 | |
| SRR4184727 | SRR29176998 | SRR3578163 | SRR30557513 | |

Table S3: Virulence factor genes (using 3 letter nomenclature) in non-O157 STEC isolates (n=286). Their presence in intimin positive isolates, enrichment factor and Jaccard similarity is also listed.

| gene (three letter code) | n_isol stx+ | n_isol stx+eae+ | n_isol stx+eae- | enrichment factor | jaccard |
| --- | --- | --- | --- | --- | --- |
| *cia* | 37 | 23 | 14 | 3.88 | 0.23 |
| *sfa* | 5 | 0 | 5 | 0 | 0 |
| *fde* | 237 | 62 | 175 | 0.84 | 0.24 |
| *usp* | 12 | 2 | 10 | 0.47 | 0.02 |
| *pap* | 16 | 1 | 15 | 0.16 | 0.01 |
| *aam* | 1 | 0 | 1 | 0 | 0 |
| *csg* | 281 | 85 | 196 | 1.03 | 0.3 |
| *csm* | 1 | 0 | 1 | 0 | 0 |
| *tib* | 9 | 0 | 9 | 0 | 0 |
| *vat* | 20 | 1 | 19 | 0.12 | 0.01 |
| *cdt* | 23 | 5 | 18 | 0.66 | 0.05 |
| *cea* | 57 | 4 | 53 | 0.18 | 0.03 |
| *eil* | 20 | 0 | 20 | 0 | 0 |
| *hra* | 46 | 4 | 42 | 0.23 | 0.03 |
| *tcp* | 2 | 0 | 2 | 0 | 0 |
| *cib* | 10 | 1 | 9 | 0.26 | 0.01 |
| *sep* | 8 | 6 | 2 | 7.09 | 0.07 |
| *tir* | 85 | 85 | 0 | inf | 1 |
| *iss* | 200 | 59 | 141 | 0.99 | 0.26 |
| *fyu* | 71 | 13 | 58 | 0.53 | 0.09 |
| *nle* | 84 | 83 | 1 | 196.27 | 0.97 |
| *etp* | 23 | 23 | 0 | inf | 0.27 |
| *foc* | 3 | 0 | 3 | 0 | 0 |
| *omp* | 217 | 80 | 137 | 1.38 | 0.36 |
| *anr* | 30 | 14 | 16 | 2.07 | 0.14 |
| *iuc* | 75 | 32 | 43 | 1.76 | 0.25 |
| *sen* | 67 | 0 | 67 | 0 | 0 |
| *F17* | 7 | 0 | 7 | 0 | 0 |
| *ygh* | 194 | 62 | 132 | 1.11 | 0.29 |
| *hha* | 18 | 11 | 7 | 3.72 | 0.12 |
| *yfc* | 25 | 12 | 13 | 2.18 | 0.12 |
| *iha* | 160 | 39 | 121 | 0.76 | 0.19 |
| *sub* | 79 | 0 | 79 | 0 | 0 |
| *ehx* | 170 | 65 | 105 | 1.46 | 0.34 |
| *csh* | 1 | 0 | 1 | 0 | 0 |
| *afa* | 17 | 2 | 15 | 0.32 | 0.02 |
| *gad* | 89 | 29 | 60 | 1.14 | 0.2 |
| *sat* | 1 | 0 | 1 | 0 | 0 |
| *mch* | 104 | 14 | 90 | 0.37 | 0.08 |
| *neu* | 22 | 4 | 18 | 0.53 | 0.04 |
| *sit* | 71 | 13 | 58 | 0.53 | 0.09 |
| *kps* | 95 | 0 | 95 | 0 | 0 |
| *irp* | 68 | 13 | 55 | 0.56 | 0.09 |
| *stx* | 286 | 85 | 201 | 1 | 0.3 |
| *tox* | 15 | 14 | 1 | 33.11 | 0.16 |
| *efa* | 45 | 44 | 1 | 104.05 | 0.51 |
| *fot* | 1 | 0 | 1 | 0 | 0 |
| *chu* | 60 | 18 | 42 | 1.01 | 0.14 |
| *aal* | 29 | 1 | 28 | 0.08 | 0.01 |
| *fim* | 6 | 0 | 6 | 0 | 0 |
| *cba* | 50 | 5 | 45 | 0.26 | 0.04 |
| *iro* | 33 | 13 | 20 | 1.54 | 0.12 |
| *shi* | 79 | 11 | 68 | 0.38 | 0.07 |
| *hly* | 155 | 57 | 98 | 1.38 | 0.31 |
| *aai* | 5 | 0 | 5 | 0 | 0 |
| *fed* | 4 | 0 | 4 | 0 | 0 |
| *dha* | 13 | 6 | 7 | 2.03 | 0.07 |
| *ets* | 7 | 3 | 4 | 1.77 | 0.03 |
| *ORF* | 5 | 0 | 5 | 0 | 0 |
| *cma* | 71 | 11 | 60 | 0.43 | 0.08 |
| *tra* | 224 | 58 | 166 | 0.83 | 0.23 |
| *nlp* | 286 | 85 | 201 | 1 | 0.3 |
| *cnf* | 17 | 1 | 16 | 0.15 | 0.01 |
| *col* | 65 | 7 | 58 | 0.29 | 0.05 |
| *yeh* | 283 | 84 | 199 | 1 | 0.3 |
| *ire* | 100 | 1 | 99 | 0.02 | 0.01 |
| *ibe* | 13 | 6 | 7 | 2.03 | 0.07 |
| *eae* | 85 | 85 | 0 | inf | 1 |
| *epe* | 19 | 3 | 16 | 0.44 | 0.03 |
| *ter* | 286 | 85 | 201 | 1 | 0.3 |
| *cva* | 34 | 13 | 21 | 1.46 | 0.12 |
| *fae* | 30 | 1 | 29 | 0.08 | 0.01 |
| *Asl* | 101 | 35 | 66 | 1.25 | 0.23 |
| *pic* | 24 | 0 | 24 | 0 | 0 |
| *lys* | 16 | 5 | 11 | 1.07 | 0.05 |
| *esp* | 133 | 85 | 48 | 4.19 | 0.64 |
| *tia* | 96 | 11 | 85 | 0.31 | 0.06 |
| *lpf* | 186 | 38 | 148 | 0.61 | 0.16 |
| *cap* | 7 | 3 | 4 | 1.77 | 0.03 |
| *cif* | 38 | 37 | 1 | 87.49 | 0.43 |
| *sig* | 2 | 0 | 2 | 0 | 0 |
| *est* | 17 | 0 | 17 | 0 | 0 |
| *iut* | 76 | 32 | 44 | 1.72 | 0.25 |
| *kat* | 33 | 23 | 10 | 5.44 | 0.24 |
| *clb* | 4 | 0 | 4 | 0 | 0 |

Table S4: Virulence factor genes (using 4 letter nomenclature) in non-O157 STEC isolates (n=286). Their presence in intimin positive isolates, enrichment factor and Jaccard similarity is also listed.

| gene (4 letter code) | n_isol stx+ | n_isol stx+eae+ | n_isol stx+eae- | enrichment | jaccard |
| --- | --- | --- | --- | --- | --- |
| *cia* | 37 | 23 | 14 | 3.88 | 0.23 |
| *faeI* | 26 | 1 | 25 | 0.09 | 0.01 |
| *hlyA* | 13 | 0 | 13 | 0 | 0 |
| *csmG* | 1 | 0 | 1 | 0 | 0 |
| *usp* | 12 | 2 | 10 | 0.47 | 0.02 |
| *papC* | 16 | 1 | 15 | 0.16 | 0.01 |
| *faeE* | 29 | 1 | 28 | 0.08 | 0.01 |
| *senB* | 67 | 0 | 67 | 0 | 0 |
| *toxB* | 15 | 14 | 1 | 33.11 | 0.16 |
| *espJ* | 45 | 44 | 1 | 104.05 | 0.51 |
| *espB* | 53 | 53 | 0 | inf | 0.62 |
| *fotE* | 1 | 0 | 1 | 0 | 0 |
| *sfaD* | 5 | 0 | 5 | 0 | 0 |
| *sitA* | 71 | 13 | 58 | 0.53 | 0.09 |
| *F17D* | 5 | 0 | 5 | 0 | 0 |
| *faeC* | 29 | 1 | 28 | 0.08 | 0.01 |
| *F17C* | 6 | 0 | 6 | 0 | 0 |
| *vat* | 20 | 1 | 19 | 0.12 | 0.01 |
| *csmA* | 1 | 0 | 1 | 0 | 0 |
| *cea* | 57 | 4 | 53 | 0.18 | 0.03 |
| *yfcV* | 25 | 12 | 13 | 2.18 | 0.12 |
| *csmD* | 1 | 0 | 1 | 0 | 0 |
| *focG* | 3 | 0 | 3 | 0 | 0 |
| *hra* | 46 | 4 | 42 | 0.23 | 0.03 |
| *faeH* | 12 | 0 | 12 | 0 | 0 |
| *nleC* | 57 | 56 | 1 | 132.42 | 0.65 |
| *cib* | 10 | 1 | 9 | 0.26 | 0.01 |
| *tir* | 85 | 85 | 0 | inf | 1 |
| *iss* | 200 | 59 | 141 | 0.99 | 0.26 |
| *neuC* | 22 | 4 | 18 | 0.53 | 0.04 |
| *eae-* | 85 | 85 | 0 | inf | 1 |
| *kpsM* | 92 | 0 | 92 | 0 | 0 |
| *ireA* | 100 | 1 | 99 | 0.02 | 0.01 |
| *F17G* | 5 | 0 | 5 | 0 | 0 |
| *ompT* | 217 | 80 | 137 | 1.38 | 0.36 |
| *tibC* | 9 | 0 | 9 | 0 | 0 |
| *sfaE* | 5 | 0 | 5 | 0 | 0 |
| *dhaK* | 13 | 6 | 7 | 2.03 | 0.07 |
| *afaB* | 13 | 2 | 11 | 0.43 | 0.02 |
| *iucC* | 75 | 32 | 43 | 1.76 | 0.25 |
| *cnf3* | 1 | 1 | 0 | inf | 0.01 |
| *papA* | 1 | 0 | 1 | 0 | 0 |
| *anr* | 30 | 14 | 16 | 2.07 | 0.14 |
| *fyuA* | 71 | 13 | 58 | 0.53 | 0.09 |
| *stx2* | 158 | 50 | 108 | 1.09 | 0.26 |
| *csgA* | 281 | 85 | 196 | 1.03 | 0.3 |
| *cshD* | 1 | 0 | 1 | 0 | 0 |
| *estb* | 3 | 0 | 3 | 0 | 0 |
| *yehA* | 258 | 83 | 175 | 1.12 | 0.32 |
| *sigA* | 2 | 0 | 2 | 0 | 0 |
| *traJ* | 59 | 24 | 35 | 1.62 | 0.2 |
| *faeF* | 21 | 1 | 20 | 0.12 | 0.01 |
| *hha* | 18 | 11 | 7 | 3.72 | 0.12 |
| *etsC* | 7 | 3 | 4 | 1.77 | 0.03 |
| *terC* | 286 | 85 | 201 | 1 | 0.3 |
| *sepA* | 8 | 6 | 2 | 7.09 | 0.07 |
| *stx1* | 176 | 41 | 135 | 0.72 | 0.19 |
| *ehxA* | 170 | 65 | 105 | 1.46 | 0.34 |
| *mchF* | 101 | 14 | 87 | 0.38 | 0.08 |
| *lpfA* | 186 | 38 | 148 | 0.61 | 0.16 |
| *F17A* | 5 | 0 | 5 | 0 | 0 |
| *iha* | 160 | 39 | 121 | 0.76 | 0.19 |
| *kpsE* | 95 | 0 | 95 | 0 | 0 |
| *fedC* | 4 | 0 | 4 | 0 | 0 |
| *fimF* | 2 | 0 | 2 | 0 | 0 |
| *fimH* | 6 | 0 | 6 | 0 | 0 |
| *espC* | 4 | 4 | 0 | inf | 0.05 |
| *nleB* | 76 | 75 | 1 | 177.35 | 0.87 |
| *gad* | 89 | 29 | 60 | 1.14 | 0.2 |
| *sat* | 1 | 0 | 1 | 0 | 0 |
| *cnf1* | 1 | 0 | 1 | 0 | 0 |
| *afaA* | 14 | 2 | 12 | 0.39 | 0.02 |
| *hlyF* | 21 | 13 | 8 | 3.84 | 0.14 |
| *aaiC* | 5 | 0 | 5 | 0 | 0 |
| *irp2* | 68 | 13 | 55 | 0.56 | 0.09 |
| *ibeA* | 13 | 6 | 7 | 2.03 | 0.07 |
| *tcpC* | 2 | 0 | 2 | 0 | 0 |
| *aamR* | 1 | 0 | 1 | 0 | 0 |
| *espY* | 26 | 12 | 14 | 2.03 | 0.12 |
| *yehD* | 277 | 82 | 195 | 0.99 | 0.29 |
| *epeA* | 19 | 3 | 16 | 0.44 | 0.03 |
| *yghJ* | 194 | 62 | 132 | 1.11 | 0.29 |
| *hlyE* | 142 | 55 | 87 | 1.49 | 0.32 |
| *eilA* | 20 | 0 | 20 | 0 | 0 |
| *faeD* | 19 | 1 | 18 | 0.13 | 0.01 |
| *esta* | 15 | 0 | 15 | 0 | 0 |
| *ORF3* | 5 | 0 | 5 | 0 | 0 |
| *espA* | 83 | 83 | 0 | inf | 0.98 |
| *cba* | 50 | 5 | 45 | 0.26 | 0.04 |
| *cdt-* | 23 | 5 | 18 | 0.66 | 0.05 |
| *shiA* | 79 | 11 | 68 | 0.38 | 0.07 |
| *AslA* | 101 | 35 | 66 | 1.25 | 0.23 |
| *espF* | 40 | 40 | 0 | inf | 0.47 |
| *aalF* | 17 | 1 | 16 | 0.15 | 0.01 |
| *fotT* | 1 | 0 | 1 | 0 | 0 |
| *katP* | 33 | 23 | 10 | 5.44 | 0.24 |
| *nleA* | 68 | 67 | 1 | 158.44 | 0.78 |
| *ORF4* | 5 | 0 | 5 | 0 | 0 |
| *capU* | 7 | 3 | 4 | 1.77 | 0.03 |
| *aalB* | 10 | 0 | 10 | 0 | 0 |
| *chuA* | 60 | 18 | 42 | 1.01 | 0.14 |
| *espP* | 49 | 34 | 15 | 5.36 | 0.34 |
| *etpD* | 23 | 23 | 0 | inf | 0.27 |
| *csmB* | 1 | 0 | 1 | 0 | 0 |
| *cma* | 71 | 11 | 60 | 0.43 | 0.08 |
| *colE* | 65 | 7 | 58 | 0.29 | 0.05 |
| *yehC* | 263 | 73 | 190 | 0.91 | 0.27 |
| *subA* | 79 | 0 | 79 | 0 | 0 |
| *efa1* | 45 | 44 | 1 | 104.05 | 0.51 |
| *nlpI* | 286 | 85 | 201 | 1 | 0.3 |
| *afaC* | 12 | 2 | 10 | 0.47 | 0.02 |
| *fotF* | 1 | 0 | 1 | 0 | 0 |
| *afaD* | 14 | 2 | 12 | 0.39 | 0.02 |
| *cvaC* | 34 | 13 | 21 | 1.46 | 0.12 |
| *fdeC* | 237 | 62 | 175 | 0.84 | 0.24 |
| *iroN* | 33 | 13 | 20 | 1.54 | 0.12 |
| *cnf2* | 15 | 0 | 15 | 0 | 0 |
| *clbB* | 4 | 0 | 4 | 0 | 0 |
| *mchC* | 82 | 12 | 70 | 0.41 | 0.08 |
| *pic* | 24 | 0 | 24 | 0 | 0 |
| *lys* | 16 | 5 | 11 | 1.07 | 0.05 |
| *traT* | 222 | 57 | 165 | 0.82 | 0.23 |
| *sfaS* | 2 | 0 | 2 | 0 | 0 |
| *tia* | 96 | 11 | 85 | 0.31 | 0.06 |
| *aalH* | 15 | 1 | 14 | 0.17 | 0.01 |
| *fotS* | 1 | 0 | 1 | 0 | 0 |
| *cif* | 38 | 37 | 1 | 87.49 | 0.43 |
| *yehB* | 260 | 75 | 185 | 0.96 | 0.28 |
| *iutA* | 76 | 32 | 44 | 1.72 | 0.25 |
| *espI* | 40 | 17 | 23 | 1.75 | 0.16 |
